# Structural insights into ligand-recognition, activation, and signaling-bias at the complement C5a receptor, C5aR1

**DOI:** 10.1101/2023.01.14.524051

**Authors:** Shirsha Saha, Jagannath Maharana, Manish K. Yadav, Parishmita Sarma, Vinay Singh, Samanwita Mohapatra, Chahat Soni, Sayantan Saha, Sudha Mishra, Manisankar Ganguly, Mohamed Chami, Ramanuj Banerjee, Arun K. Shukla

## Abstract

Activation of the complement cascade is a critical part of our innate immune response against invading pathogens, and it operates in a concerted fashion with the antibodies and phagocytic cells towards the clearance of pathogens. The complement peptide C5a, generated during the activation of complement cascade, is a potent inflammatory molecule, and increased levels of C5a are implicated in multiple inflammatory disorders including the advanced stages of COVID-19 pathophysiology. The proximal step in C5a-mediated cellular and physiological responses is its interaction with two different seven transmembrane receptors (7TMRs) known as C5aR1 and C5aR2. Despite a large body of functional data on C5a-C5aR1 interaction, direct visualization of their interaction at high-resolution is still lacking, and it represents a significant knowledge gap in our current understanding of complement receptor activation and signaling. Here, we present cryo-EM structures of C5aR1 activated by its natural agonist C5a, and a G-protein-biased synthetic peptide ligand C5a^pep^, in complex with heterotrimeric G-proteins. The C5a-C5aR1 structure reveals the ligand binding interface involving the N-terminus and extracellular loops of the receptor, and we observe that C5a exhibits a significant conformational change upon its interaction with the receptor compared to the basal conformation. On the other hand, the structural details of C5a^pep^-C5aR1 complex provide a molecular basis to rationalize the ability of peptides, designed based on the carboxyl-terminus sequence of C5a, to act as potent agonists of the receptor, and also the mechanism underlying their biased agonism. In addition, these structural snapshots also reveal activation-associated conformational changes in C5aR1 including outward movement of TM6 and a dramatic rotation of helix 8, and the interaction interface for G-protein-coupling. In summary, this study provides previously lacking molecular basis for the complement C5a recognition and activation of C5aR1, and it should facilitate structure-based discovery of novel lead molecules to target C5aR1 in inflammatory disorders.

## Introduction

The complement system, also known as the complement cascade, is an integral part of our immune response against pathogenic infections^1^. Once activated, it plays a vital role in efficient destruction and clearance of microbial agents through the formation of membrane attack complex and associated mechanisms^2,3^. Complement activation results in the generation of several peptide fragments by the action of different proteases, and these complement peptides subsequently exert their functions through the corresponding receptors and effectors^2,3^. One such complement peptide is C5a, which is generated through the proteolytic cleavage of the complement component C5 by the C5 convertase enzyme, and it consists of 74 amino acids^2,3^. C5a is a highly potent inflammatory molecule, and its abnormal production often contributes to the onset and progression of multiple inflammatory conditions including sepsis and the advanced stage of COVID-19 pathophysiology^4-6^.

C5a binds to, and activates, two distinct seven-transmembrane receptors (7TMRs), namely the C5aR1 and C5aR2 (also known as C5L2)^7,8^. While C5aR1 is a prototypical G-protein-coupled receptor (GPCR), C5aR2 couples only to β-arrestins (βarrs) without any measurable G-protein activation, and hence it is also referred to as an Arrestin-Coupled Receptor (ACR)^8,9^. C5aR1 is widely distributed in immune cells including the macrophages and neutrophils, and endothelial cells with primary coupling to Gαi sub-type of heterotrimeric G-proteins^10^ (Figure 1A). Upon activation by agonists, C5aR1 also undergoes phosphorylation followed by the binding of βarrs and subsequent internalization^11^. The interaction of C5a with C5aR1 and ensuing downstream signaling responses have been implicated in the disease severity of COVID-19 patients, including a potential chemoattracting role that leads to infiltration of neutrophils and monocytes in the broncho-alveolar lavage fluid (BALF) of the patients^4^. Moreover, a monoclonal antibody capable of blocking C5a-C5aR1 interaction has shown therapeutic promise for COVID-19 in mouse model^4^.

**Figure 1.**
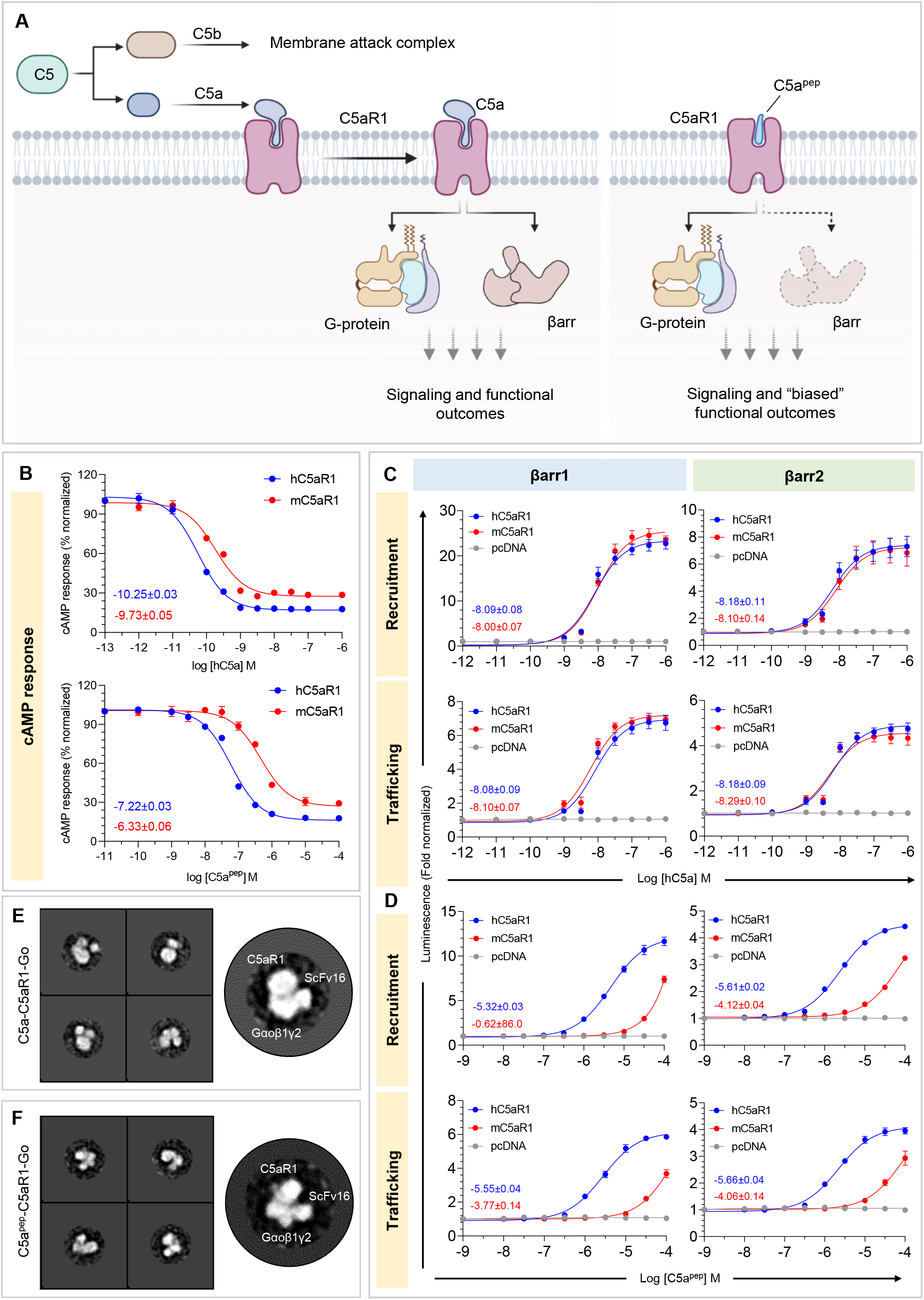
Agonist-induced activation and pharmacology of C5aR1. **(A)** C5 convertase, a critical player in the complement cascade cleaves complement C5 into two different fragments, C5a and C5b. C5b is directed towards the formation of the pathogen killing membrane attack complex (MAC), whereas C5a activates the cognate GPCR, C5aR1. C5aR1 is a classical G protein-coupled receptor and couples to Gαi subtype of heterotrimeric G-proteins and β-arrestins upon stimulation with C5a resulting in various cellular responses. C5aR1 also recognizes a C5a-derived peptide agonist, C5a^pep^ which drives signaling through G-proteins whereas weakly recruits βarrs. C5a^pep^ triggers “biased” functional outcomes upon binding to C5aR1. Schematic prepared using BioRender.com. **(B)** C5a (top) and C5a^pep^ (bottom) driven Gαi-mediated second messenger response as measured by agonist dependent decrease in forskolin-induced cytosolic cAMP levels downstream to C5aR1. Respective logEC50 values are mentioned in the inset. Data (mean±SEM) represents four independent experiments, normalized with respect to highest signal (measured as 100%) for each receptor. **(C-D)** C5a/C5a^pep^ induced βarr1/2 recruitment and trafficking as measured by NanoBit assay. Respective logEC50 values are mentioned in the inset. Data (mean±SEM) represents four independent experiments, fold normalized with respect to luminescence observed at lowest dose (measured as 1) for each receptor. **(E, F)** Visualization of the purified C5a-C5aR1-Go (top) and C5a^pep^-C5aR1-Go (bottom) complexes via negative staining EM. 2D class averages and representative 2D class depicting a typical GPCR-G-protein complex are shown.

Previously determined crystal structures of C5a have revealed a four-helix architecture with connecting loops stabilized by three disulfide bridges^12^. A series of mutagenesis studies, coupled with functional assays, have suggested that binding of C5a with C5aR1 involves an interface between the core region of C5a with the N-terminus and extracellular loop 2 (ECL2) of C5aR1, and a second interface between the carboxyl-terminus of C5a with the extracellular pocket of the receptor^13-16^. In addition to C5a, several peptide ligands derived from, and modified based on the carboxyl-terminus sequence of C5a, have been described as potent agonists for C5aR1, albeit with relatively lower affinities^16-21^. Of these, a hexapeptide referred to as C5a^pep^ is particularly interesting as it behaves as a functionally-biased agonist compared to C5a in terms of transducer-coupling and cellular responses^11^. It exhibits comparable efficacy to C5a for cAMP inhibition and ERK1/2 MAP kinase activation, although with significantly weaker potency, while it is a partial agonist for βarr recruitment and trafficking^11^. Moreover, C5a^pep^ displays full agonism for inhibiting LPS-induced IL-6 release in human macrophages but only partial agonism with respect to neutrophil migration^11^. However, direct visualization of agonist-binding to the receptor, either with C5a or peptide agonists, is currently lacking, and it remains a major knowledge gap in our understanding of complement recognition mechanism and activation of C5aR1.

In this study, we present two cryo-EM structures of C5aR1 in complex with the heterotrimeric G-proteins where the receptor is occupied either by C5a or C5a^pep^. These structures not only unravel the molecular basis of complement recognition and receptor activation including a previously unanticipated dramatic rotation of helix 8, but they also offer important insights into functional bias at the receptor elicited by peptide agonists. Additionally, these structural snapshots also offer a previously lacking platform to facilitate structure-guided novel ligand discovery at the complement receptors with enhanced sub-type selectivity and improved biased agonism.

## Results and discussion

### Reconstitution of agonist-C5aR1-G-protein complexes

In order to determine the structure of active C5aR1 in complex with G-proteins, we first started with expression and purification of the full-length human C5aR1 using the baculovirus expression system. However, despite robust expression and efficient purification, the receptor exhibited a heterogenous profile when analyzed by size-exclusion chromatography. Therefore, we focused our efforts on the mouse C5aR1, which appeared highly monodisperse and suitable for structural studies (Figure S1A). We first compared the pharmacology of human C5a (hC5a) and C5a^pep^ on human and mouse C5aR1, referred to as hC5aR1 and mC5aR1, respectively, in terms of G-protein-coupling, and βarr recruitment and trafficking using previously described GloSensor^22^ and NanoBiT^23^ assays, respectively. We observed that both hC5a and C5a^pep^ behave as full agonists on mC5aR1 with slightly lower potency compared to hC5aR1 in terms of G-protein-mediated cAMP response (Figure 1B). On the other hand, while hC5a exhibits full agonism for βarr recruitment and endosomal trafficking on mC5aR1 (Figure 1C), C5a^pep^ displays partial agonism at mC5aR1 compared to hC5aR1 in these assays (Figure 1D), and it exhibits G-protein-bias in line with our previous study^11^. In these experiments, mC5aR1 and hC5aR1 were expressed at comparable levels (Figure S1B-K). Subsequently, we successfully reconstituted C5a-C5aR1-Gαoβ1γ2 and C5a^pep^-C5aR1-Gαoβ1γ2 complexes stabilized using ScFv16 by combining the purified components (Figure S2A-B), and negative-staining of these complexes suggested uniform particle distribution with an overall complex architecture reminiscent of typical GPCR-G-protein assemblies (Figure 1E-F and S2C-D).

### Overall structures of C5a/C5a^pep^-C5aR1-Gαoβ1γ2 complexes

These complexes were subsequently subjected to cryo-EM data collection on a 300kV Titan Krios microscope followed by data analysis using cryoSPARC (v3.3.2/v4)^24^ as outlined in Figure S3 for the C5a-C5aR1-Gαoβ1γ2-ScFv16 and Figure S4 for the C5a^pep^-C5aR1-Gαoβ1γ2-ScFv16 complex yielding structures at 3.9Å and 3.4Å, respectively (Figure 2 and S5). Despite somewhat moderate resolution, cryo-EM maps allowed unambiguous modeling of the secondary structures of all the components including C5a and C5a^pep^ in the corresponding structures (Figure 2 and Figure S6A-B). The final model of C5a-C5aR1-Go complex contains clear density for the residues ranging from Pro24^N-term^ to Ser315^H8^ of the receptor, although the residues Tyr103 to Asp105 in ECL1 and Val187 to Glu200 in ECL2 did not show discernible densities, potentially due to inherent flexibility (Figure S7). The C5a^pep^-C5aR1-Go complex contains clear densities for the residues ranging from Gly36^N-term^ to Ser315^H8^ although the regions corresponding to Ala66 to Arg68 in ICL1 and Lys179 to Glu200 in ECL2 were not resolved in the final model (Figure S7). A schematic representation of the residues corresponding to the different components of the complex resolved in the final structural models are summarized in Figure S7. Expectedly, the overall quality of cryo-EM maps was the highest at the receptor-Go interface while the extracellular loops exhibited relatively higher variability. The overall structures of C5a^pep^ and C5a-bound C5aR1 are similar and exhibit an RMSD of 0.736Å across the receptor upon superimposition (Figure S8).

**Figure 2.**
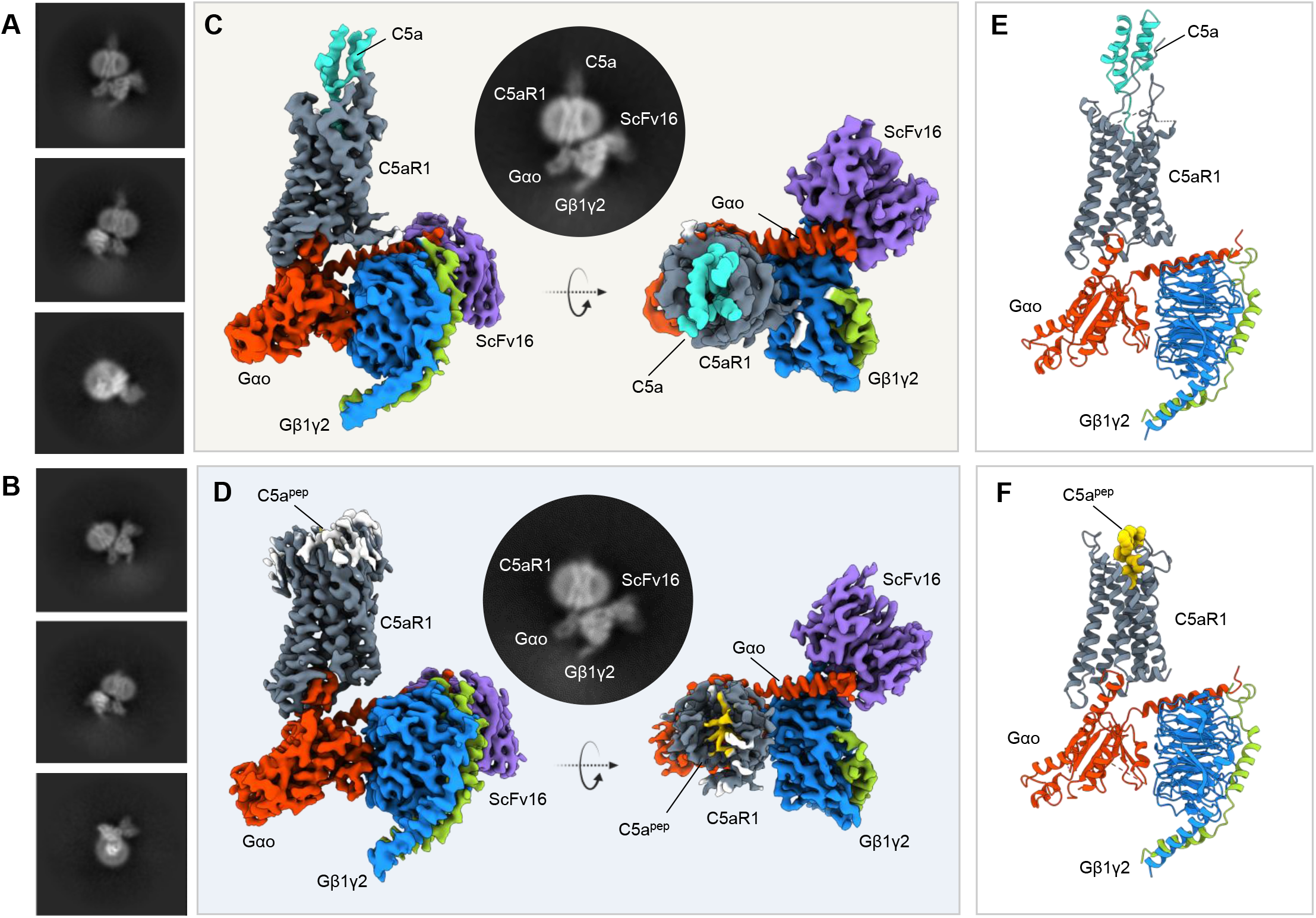
Overall structure of C5a- and C5apep-bound C5aR1-G-protein complexes. **(A, B)** Cryo-EM 2D class averages of C5a bound C5aR1-G-protein complex and C5a^pep^-C5aR1-G-protein complex respectively. **(C, D)** Two different views showing subunit organization of the C5a and C5a^pep^ bound C5aR1-G-protein complexes respectively; representative 2D class average of the complex with clear secondary features have been included in inset. **(E)** Ribbon diagram of the C5a bound C5aR1 complex (gray: C5aR1, cyan: C5a, orange: Gαo, blue: Gβ1, green: Gɣ2). **(F)** Ribbon diagram of the C5a^pep^ bound C5aR1 complex (gray: C5aR1, yellow: C5a^pep^, orange: Gαo, blue: Gβ1, green: Gɣ2, purple: ScFv16).

### Interaction of C5a and C5a^pep^ with C5aR1

Previous crystal structures of C5a have revealed a rigid core consisting of a four-helix bundle (H1-H4) wherein the carboxyl-terminus adopts a short α-helical conformation^12^ (Figure 3A). Strikingly, we observed a significant conformational and structural rearrangement in C5a upon its interaction with C5aR1, although it still adopts a four-helix bundle architecture (Figure 3B-C). In particular, the carboxyl-terminus of C5a displays an extended conformation instead of the short α-helical turn, and the third short helix tilted at an angle of about 45º in C5aR1-bound conformation compared to the basal state (Figure 3C). Consistent with previous studies, we observed a two-site binding mechanism of C5a to C5aR1 (Figure 3D). The N-terminus and ECL2 of the receptor interface with the central core of C5a, while the extended carboxyl-terminus of C5a docks into the orthosteric binding pocket formed on the extracellular side of the transmembrane bundle of the receptor (Figure 3D). The interface areas of these two sites of C5a interaction on C5aR1 are 316Å^2^ and 600Å^2^, respectively. There are several hydrogen bonds, hydrophobic interactions, salt bridges, and polar interactions that help stabilize the overall positioning of C5a on C5aR1 (Figure 3E and S9A). For example, His29 in the N-terminus of C5aR1 forms a hydrogen bond and non-bonded contact with Arg^37^ and Arg^40^ of C5a, respectively. In addition, non-bonded contacts between Ile28 in the N-terminus of C5aR1 with Arg^40^ and Ile^41^, and hydrogen bonds between Glu176 in ECL2 of the receptor and His^67^ and Ser^66^ of C5a are also key determinants for the first binding site. On the other hand, as a part of the second binding site, Gly^73^ and Arg^74^ in C5a form extensive interactions with Val287^7.38^ and Tyr259^6.51^, Gly263^6.55^ and Ile266^6.58^, respectively, in TM6 of the receptor (Figure 3D-E).

**Figure 3.**
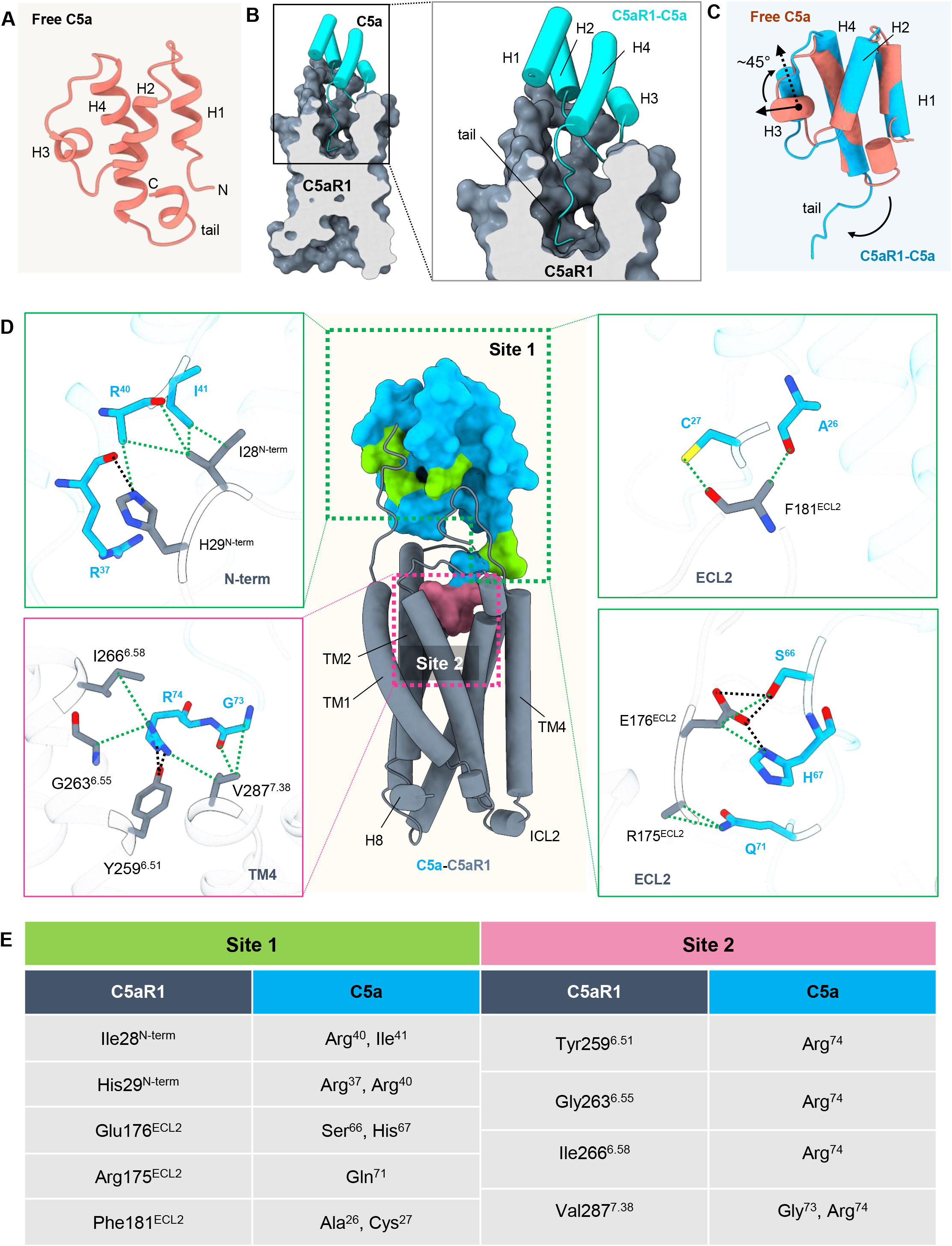
Structural details of C5a-C5aR1 interaction. **(A)** Overall architecture of free C5a showing four helical bundle and a short helix in the C tail. **(B)** Upon binding to C5aR1, the C-terminal tail of C5a docks perpendicularly into the ligand binding cavity. **(C)** Structural comparison of free C5a with C5a bound to C5aR1. Helix 3 of C5a can be seen to exhibit a rotation of ∼45º upon binding to the receptor. **(D)** Interaction interfaces of site 1 and site 2 of C5a with the N-terminus, ECL2 and TMs of C5aR1 have been illustrated. **(E)** A comprehensive list of all the interactions including polar and non-bonded contacts have been included in the table. (Polar interactions: Black dotted lines, Non-bonded contacts: Green dotted lines).

In the C5a^pep^-C5aR1-Go structure, we observe that C5a^pep^ adopts a peg-like conformation and positions itself into the orthosteric pocket with an interface that is analogous to the carboxyl-terminus of C5a (Figure 4B). Interestingly, the N-methyl-phenylalanine residue of C5a^pep^ is located in close vicinity of ECL2 and forms an anion-π interaction with Glu176 and non-bonded contact with Tyr178 in the ECL2 of the receptor (Figure 4A, C). In addition, Pro^3^ of C5a^pep^ makes a contact with Glu280^7.32^ in the receptor through the main chain oxygen atom, while Cha^5^ of C5a^pep^ forms hydrogen bonds with Arg175^ECL2^ in the receptor (Figure 4C). Finally, d-Arg^6^ in C5a^pep^ makes substantial interactions with the residues in TM2, TM3 and TM7 of the receptor (Figure 4C). Interestingly, d-Cha^4^ in C5a^pep^ does not appear to be involved in any major interaction with the receptor. These extensive interactions help stabilize C5a^pep^ in the receptor’s orthosteric pocket, and a comprehensive map of interactions between C5a/C5a^pep^ and C5aR1 are listed in Figure S9.

**Figure 4.**
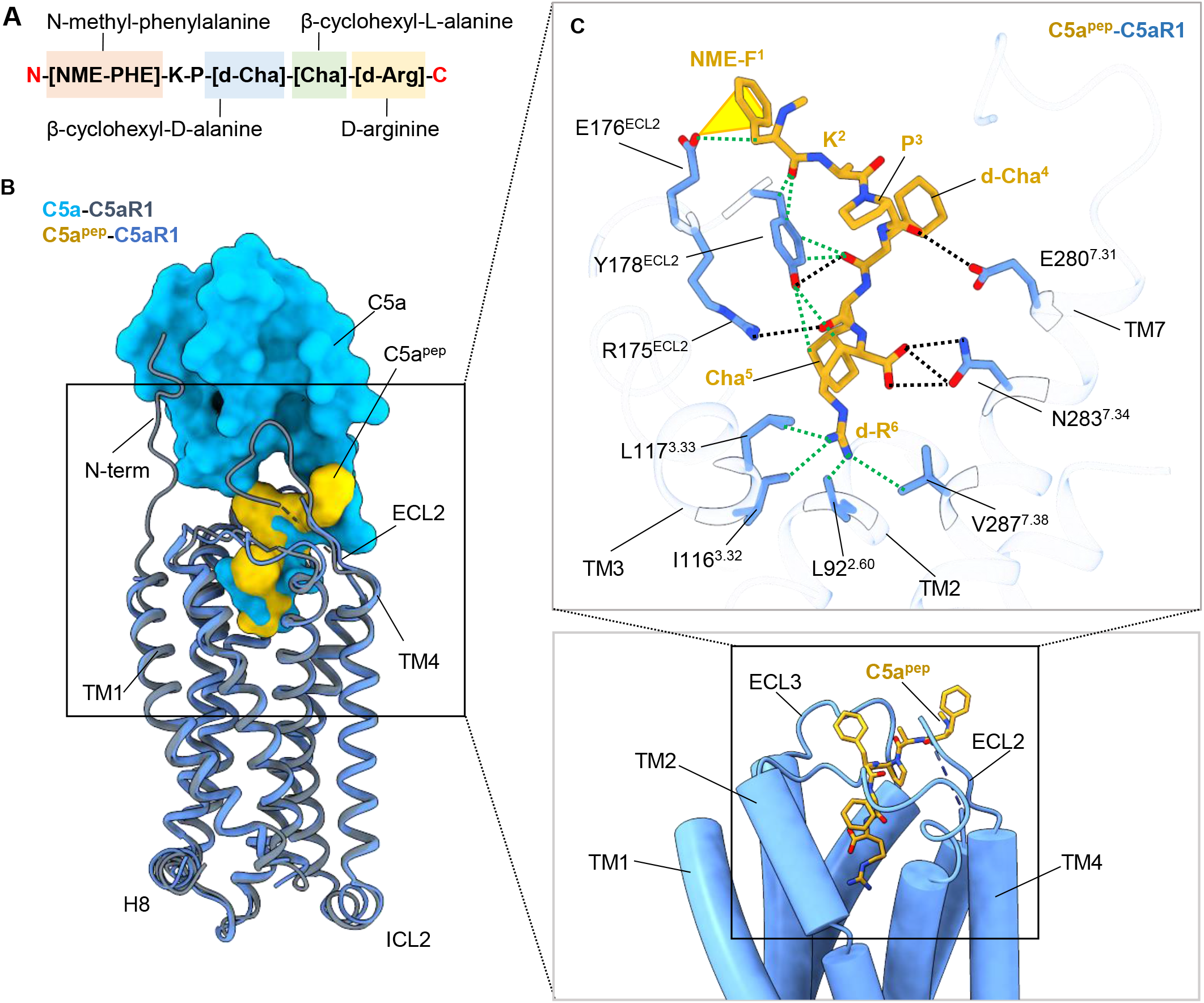
Structural details of C5apep-C5aR1 interaction. **(A)** Sequence of the C5a derived peptide, C5a^pep^. **(B)** C5a^pep^ occupies a similar binding pocket as C5a in C5aR1. **(C)** The binding pocket of C5a^pep^ in C5aR1 on the extracellular side is surrounded by residues from ECLs and TMs. Interactions of the residues in C5a^pep^ with C5aR1 in the ligand binding pocket have been represented as dotted lines. The yellow cone depicts anion-π interaction between NME-F1 of C5a^pep^ and Glu176^ECL2^ of C5aR1. (Polar interactions: Black dotted lines, Non-bonded contacts: Green dotted lines).

A direct comparison of the second C5a binding site on C5aR1 with that of C5a^pep^ binding site reveals that their engagement with Leu92^2.60^, Arg175^ECL2^ and Glu176^ECL2^, and Val287^7.38^ are common (Figure 5A-B). However, there are substantial differences in the binding modes of the two ligands and the receptor residues engaged by them (Figure 5C). The carboxyl-terminus of C5a adopts a hook-like conformation and penetrates deeper into the orthosteric pocket as compared to C5a^pep^ (Figure 5A). Leu^72^ in C5a engages with Thr95^2.63^ and Asn100^ECL1^ of C5aR1 while Gln^71^ in C5a contacts Pro113^3.29^ of C5aR1. Furthermore, Gly^73^ in C5a interacts with Met120^3.36^ of the receptor through its main chain oxygen (Figure 5C). Additionally, upon binding of C5a, Glu176 in ECL2 of C5aR1 forms hydrogen bonds with Ser^66^ and His^67^ in the extended carboxyl-terminus of C5a, and these potentially serve to bridge the ECL2 of the receptor with the C-terminus of the natural agonist, together with the additional contacts mentioned above. In contrast, although the main chain oxygen of d-Cha^4^ and Cha^5^ in C5a^pep^ interact with Tyr178^ECL2^ and Arg175^ECL2^ of the receptor, respectively, the rest of the contacts with ECL2 are absent (Figure 5B-C). Instead, C5a^pep^ engages Ile116^3.32^, Leu117^3.33^, Tyr178^ECL2^, Glu280^7.31^ and Asn283^7.34^, which are absent in the case of C5a (Figure 5C). Finally, Arg^74^ in C5a engages with Gly263^6.55^ and Ile266^6.58^ in C5aR1 through hydrophobic interactions, and it also makes ionic contact with Tyr259^6.51^. In contrast to this, d-Arg^6^ in C5a^pep^ is positioned upwards compared to Arg^74^ in C5a, and therefore, interacts instead with Ile116^3.32^ and Leu117^3.33^ of the receptor (Figure 5C and S10A).

**Figure 5.**
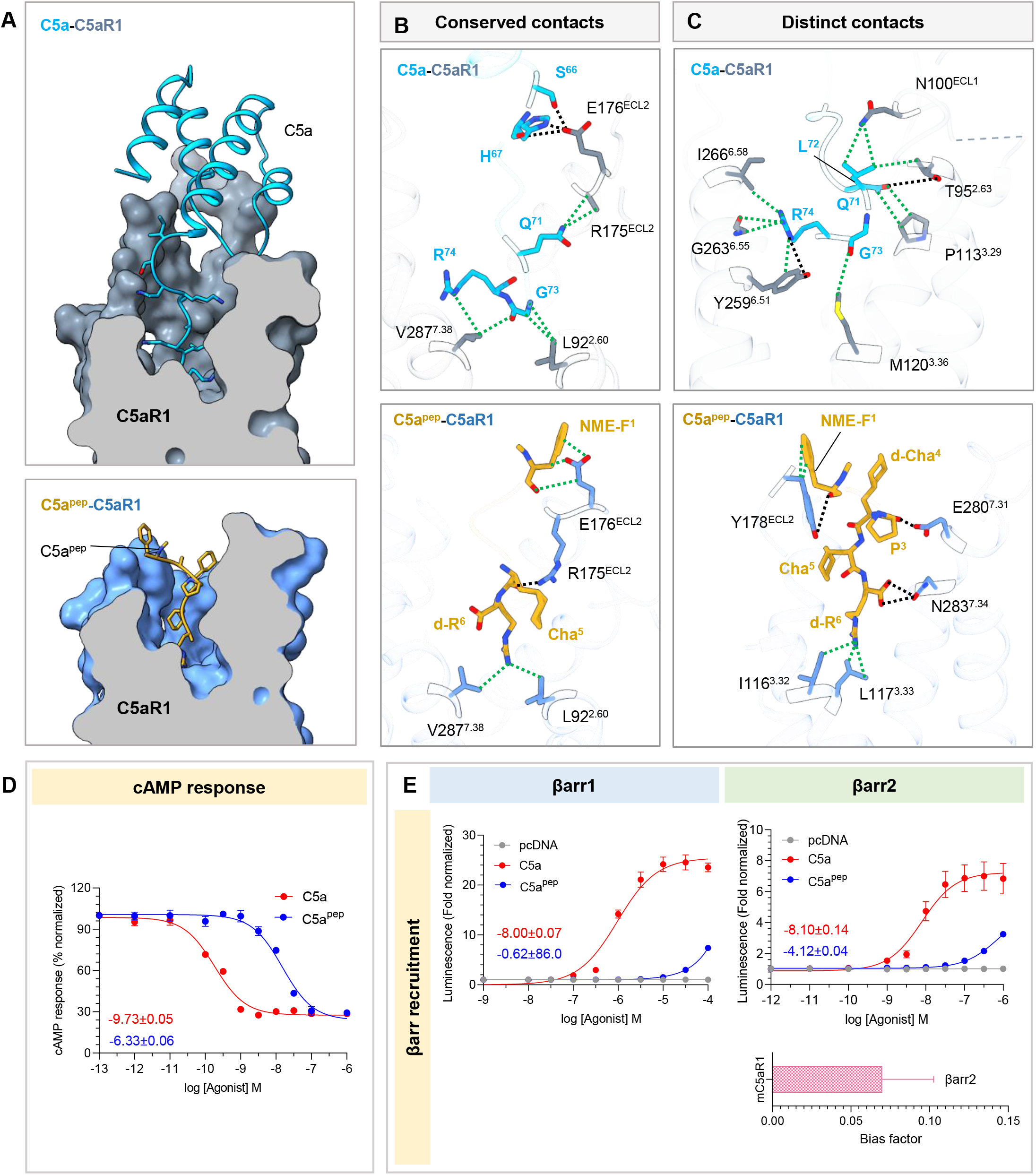
Comparison of C5a and C5a^pep^ binding to C5aR1. **(A)** The C terminal region of C5a (top) and C5a^pep^ (bottom) employ a similar binding pocket on C5aR1. The receptor and ligands have been represented as surface slices and ribbons respectively. **(B, C)** Common and unique interactions of C5a (top) and C5a^pep^ (bottom) at the ligand-receptor interface have been shown as dotted lines. (Polar interactions: Black dotted lines, Non-bonded contacts: Green dotted lines). **(D)** Comparison of C5a/C5a^pep^ mediated cAMP response downstream of mouse C5aR1 reveal reduced efficacy of C5a^pep^ as compared to C5a. Respective logEC50 values are mentioned in the inset. Data (mean±SEM) represents four independent experiments, normalized with respect to highest signal (measured as 100%) in response to each ligand. **(E)** Measuring βarr1/2 recruitment to mouse C5aR1 upon stimulation with C5a and C5a^pep^ shows significant reduction in both efficacy as well as potency of C5a^pep^ as compared to C5a (top). Respective logEC50 values are mentioned in the inset. Data (mean±SEM) represents four independent experiments, fold normalized with respect to luminescence observed at lowest dose (measured as 1) for each ligand. Bias factor (β value) determined taking C5a as reference elucidates the G-protein biased nature of C5a^pep^. The graphs in panel D and E are derived from the same experimental data that are presented in Figure 1B-D.

It is conceivable that the lack of first binding site in case of C5a^pep^ would impart lower binding affinity to the receptor compared to C5a as proposed based on the two-site binding mechanism^13^. However, it is also possible that the differences observed in the second binding site for these two ligands may also be responsible, at least partly, for the differences in their binding affinities to the receptor. More importantly, these distinct set of interactions formed by C5a vs. C5a^pep^ with the receptor are likely to be the primary determinants for the difference in their transducer-coupling efficacy, especially βarr recruitment, and the resulting G-protein-bias of C5a^pep^ (Figure 5D-E) although future studies are required to explore it further.

Interestingly, previous studies have suggested that the removal of terminal arginine (Arg^74^) in C5a *in-vivo* by the action of a carboxypeptidase yields C5a^des-Arg^, and it displays significantly reduced binding affinity and potency at C5aR1 when tested using recombinant ligand^10,16,25,26^. The extensive interaction of Arg^74^ in C5a with multiple residues in C5aR1 such as Tyr259^6.51^, Gly263^6.55^, Ile266^6.58^ and Val287^7.38^, which are critical for stabilizing carboxyl-terminal conformation of C5a in the orthosteric pocket, and will be absent in case of C5a^des-Arg^, may help rationalize its lower potency at the receptor (Figure S10B).

### Structural insights into species-specific differences in agonist pharmacology

As presented in Figure 1, we observed only a small difference between the human and mouse C5aR1 for C5a-induced G-protein-coupling as measured using GloSensor assay, whereas βarr interaction and trafficking were essentially identical. On the other hand, βarr recruitment and trafficking were dramatically different between the human and mouse receptors upon C5a^pep^ stimulation. Sequence analysis of the human and mouse C5aR1 in terms of C5a- and C5a^pep^-interacting residues provides the potential structural basis for this observation (Figure S11). While there are differences between C5a-interacting residues between the human and mouse receptor, especially in the second binding site, they appear to be rather modest. On the other hand, the differences are more pronounced in case of C5a^pep^ (Figure 6A and S12A). For example, Thr95^2.63^, Asn100^ECL1^, Glu176^ECL2^ and Phe181^ECL2^, and Ile266^6.58^ in mouse C5aR1 are substituted with Ser95^2.63^, His100^ECL1^, Val176^ECL2^ and Tyr181^ECL2^, and Met265^6.58^, respectively, in human C5aR1. Glu176^ECL2^ in the ECL2 of mouse C5aR1 forms hydrogen bonds with Ser^66^ and His^67^ of C5a, thereby rigidifying the conformation of C5a within the extracellular binding pocket of C5aR1. Substitution of Glu176^ECL2^ in mouse C5aR1 with Val would result in disruption of these hydrogen bonds and thereby, facilitate formation of weak hydrophobic interactions between Val and its neighboring residues, increasing the flexibility of the bound ligand. Similarly, substitution of Phe181^ECL2^ and Ile266^6.58^ in mouse C5aR1 with Tyr and Met respectively would facilitate formation of polar interactions and hydrogen bond of these residues with their surrounding environment. These substitutions in C5aR1 might provide a plausible explanation for the small difference observed in the GloSensor assay. On the other hand, C5a^pep^-interacting residues Tyr178 ^ECL2^, and Glu280^7.31^ and Asn283^7.34^ in mouse C5aR1 are substituted with Arg178 ^ECL2^, Lys279^7.31^ and Asp282^7.34^, respectively, in human C5aR1 (Figure 6B and S12B). In the structure, Tyr178^ECL2^ interacts with NME-Phe^1^, d-Cha^4^ and Cha^5^, and substitution of Tyr with Arg would reverse the polarity in these positions. Likewise, substitution of Glu280^7.31^ and Asn283^7.34^ in mouse C5aR1 with Lys and Asp respectively would alter the individual polarity patterns and possibly allow differential interactions in these sites. These alterations between the amino acid sequence of mouse and human C5aR1 might account for the difference observed in βarr recruitment and trafficking between the human and mouse receptors upon C5a^pep^ stimulation. In fact, Tyr178Arg mutation in mouse C5aR1 enhances the potency and efficacy of C5a^pep^ in βarr1 trafficking, which supports the structural interpretation as outlined above (Figure 6C). While additional studies are required to link this primary sequence difference with the observed responses in the functional assays, it provides a plausible structural explanation for species-specific differences in ligand potency for C5aR1 ligands as reported previously^27^.

**Figure 6.**
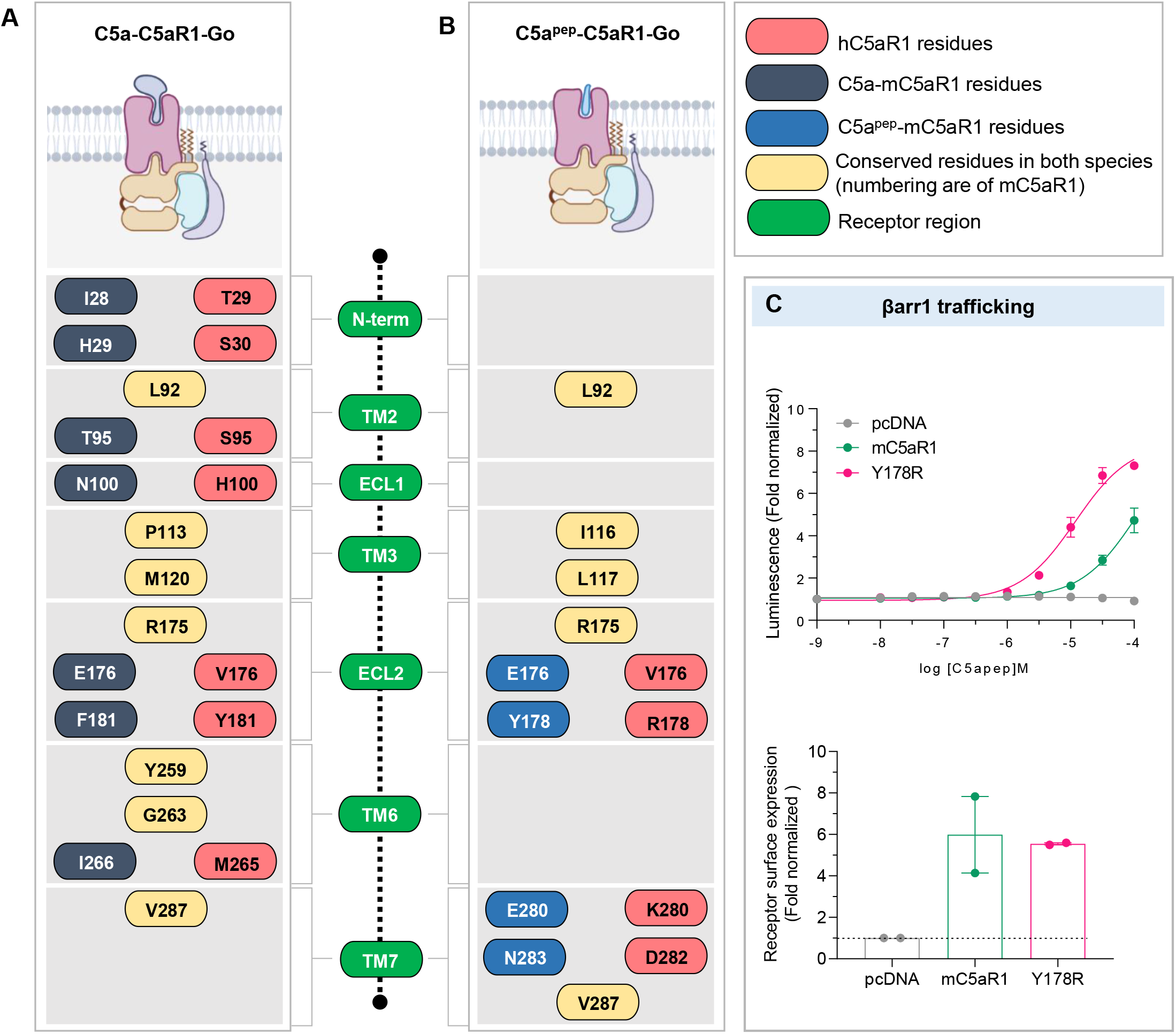
Structural insights into species-specific ligand bias at mouse C5aR1. **(A, B)** Schematic representation of residue contacts between C5a and C5a^pep^ with C5aR1. The nature of contacts annotated are highlighted in an inset box on the right. **(C)** Measuring βarr1 trafficking in response to C5a^pep^ downstream to a series of mouse C5aR1 mutants mimicking the corresponding human C5aR1 residues show dramatic increase in both potency and efficacy of βarr1 trafficking compared to the wild type mouse receptor. Data (mean±SEM) represents two independent experiments, performed in duplicate, fold normalized with respect to luminescence observed at lowest dose (measured as 1) for each receptor (top). All the receptors were expressed at comparable levels (bottom).

### Structural insights into competitive antagonism of PMX53

Finally, these structures also provide interesting clues about the competitive antagonistic behavior of some peptide fragments, PMX53 in particular, designed and modified based on the carboxyl-terminus of C5a^28-30^. The overall binding pocket of PMX53 observed in previously determined crystal structure is analogous to that of C5a^pep^ and the extended carboxyl-terminus of C5a (Figure 7A-B). The comparison of PMX53 binding with that of C5a^pep^ and the carboxyl-terminus of C5a reveals common interactions with Arg175 in ECL2, Leu92 in TM2, Glu176 in ECL2 and Val287 in TM7 of the receptor (Figure 7C). These interactions may explain the competitive binding mode of PMX53 with C5a and C5a^pep^. On the other hand, PMX53 also makes several critical contacts with C5aR1 that are absent in the C5a/C5a^pep^ bound C5aR1 structures (Figure S13). For example, previous studies have proposed that Ile116^3.32^ and Val286^7.38^ (Val287^7.38^ in case of mouse C5aR1) form an “activation switch” in C5aR1^31^, and mutating these residues completely abolishes C5a receptor activity^32^. In our structure, C5a^pep^ makes extensive contacts with both Ile116^3.32^ and Val287^7.38^, which further stabilizes the conformation of the peptide within the ligand binding pocket of the receptor. In contrast, although PMX53 makes similar contacts, it also engages several extra residues on C5aR1, that are not observed in C5a/C5a^pep^ bound structures (Figure S13). The bulky nature of Trp at position 5 of the cyclic hexapeptide possibly restricts adequate movement of I116^3.32^, thereby locking the inactive state and hindering receptor activation (Figure S13). Moreover, the cyclic nature of PMX53 blocks the C-terminal carboxylate which has been proposed to help induce agonist activity of the peptide (Figure 7D). Interestingly, it has been observed that mutating the fifth residue in C5a^pep^ to tryptophan converts it into an antagonist and substituting I116^3.32^ of C5aR1 with Ala rescues this antagonism^21,33^. Moreover, the conformation of PMX53 is stabilized by a network of hydrogen bonds formed between the residues of PMX53 with Pro113^3.29^, R175^ECL2^, Cys188^ECL2^, V190^ECL2^, Tyr258^6.51^, Thr261^6.54^ and Asp282^7.34^ of C5aR1 (Figure 7E), suggesting a rigid binding mode of PMX53 within the extracellular binding pocket of C5aR1, in turn stabilizing the receptor in inactive conformation. Taken together, these structural insights help rationalize the antagonistic nature of PMX53 despite sharing an overall similar binding pocket on the receptor as C5a/C5a^pep^.

**Figure 7.**
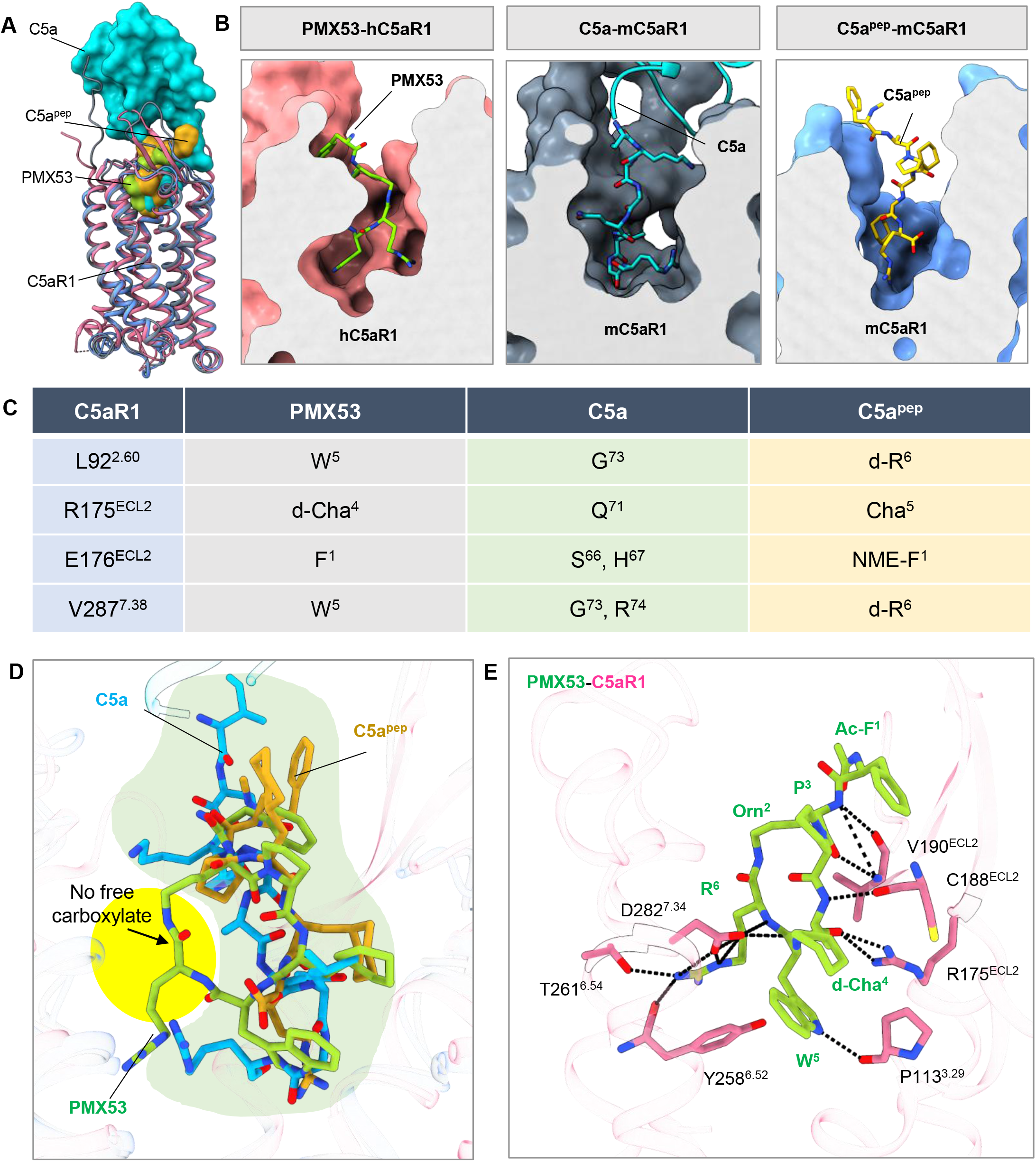
Structural insights into competitive antagonism of PMX53 at C5aR1. **(A)** Superimposition of active C5a and C5a^pep^ activated C5aR1 with the inactive PMX53 bound C5aR1 (PDB ID: 6C1R). Ligands are shown in surface and receptors in ribbon representation. **(B)** PMX53 binds at a similar pocket as C5a and C5a^pep^. Surface slices of C5aR1 with cognate ligands are depicted to highlight the occupancy of ligands at the same binding pocket. **(C)** Overall conserved interactions between PMX53, C5a and C5a^pep^ with C5aR1 are listed. **(D)** The cyclic peptide, PMX53 engages an extra binding site (yellow patch) on C5aR1 unlike C5a/C5a^pep^ (green patch). The carboxylate group is blocked in PMX53 (highlighted in yellow) due to the cyclic nature of the peptide, further preventing agonistic behavior. **(E)** PMX53 forms extensive hydrogen bonds with the residues of the ligand binding pocket of C5aR1.

### Interaction of C5aR1 with G-proteins

The overall interface between C5aR1 and G-protein heterotrimer displays an approximate buried surface area of 1,880Å^2^ and 1,720Å^2^ in C5a-C5aR1-Go and C5a^pep^-C5aR1-Go structures, respectively, with Gαo constituting the primary interface (Figure 8A-B). Expectedly, the distal end of the α5-helix of Gαo inserts into the cytoplasmic side of the receptor transmembrane core as observed for other GPCR-G-protein complexes including a close phylogenetic neighbor of C5aR1, the Formyl Peptide receptor subtype 2 (FPR2)^34-36^ (Figure 8C-E). On the receptor side, the major interface is formed by the cytoplasmic ends of TM2, TM3, TM6, TM7, ICL2 and ICL3, and is essentially identical between C5a and C5a^pep^-bound structures (Figure S14). The overall interaction between the receptor and Gαo is stabilized by multiple hydrogen bonds, polar contacts, and hydrophobic interactions, which are listed in Figure S14. Some of the critical interactions include hydrogen bonding between Asn^347^, Asp^341^ and Tyr^354^ of the α5 helix in Gαo with Arg148^ICL2^, Arg233^5.68^ and Ser238^ICL3^ of C5aR1, respectively (Figure 8C-D and S14). Interestingly, the residues ranging from Trp143 to Lys146 in the ICL2 of the receptor adopt a short α-helical turn that is positioned into the hydrophobic groove formed by the α5, αN, β1 and β3 strands of Gαo. Specifically, Ile142 in the ICL2 of C5aR1 interacts with Asn^194^ and Leu^195^ of the β2-β3 loop of Gαo, while Gln145 and Lys146 of ICL2 engage with Lys^32^ in the αN helix-β1 loop of Gαo. In addition, the receptor-G-protein engagement is facilitated further by the interaction of ICL3 residues such as Thr236, Arg237 and Ser238 with Tyr^354^ in the α5 helix of Gαo (Figure S14).

**Figure 8.**
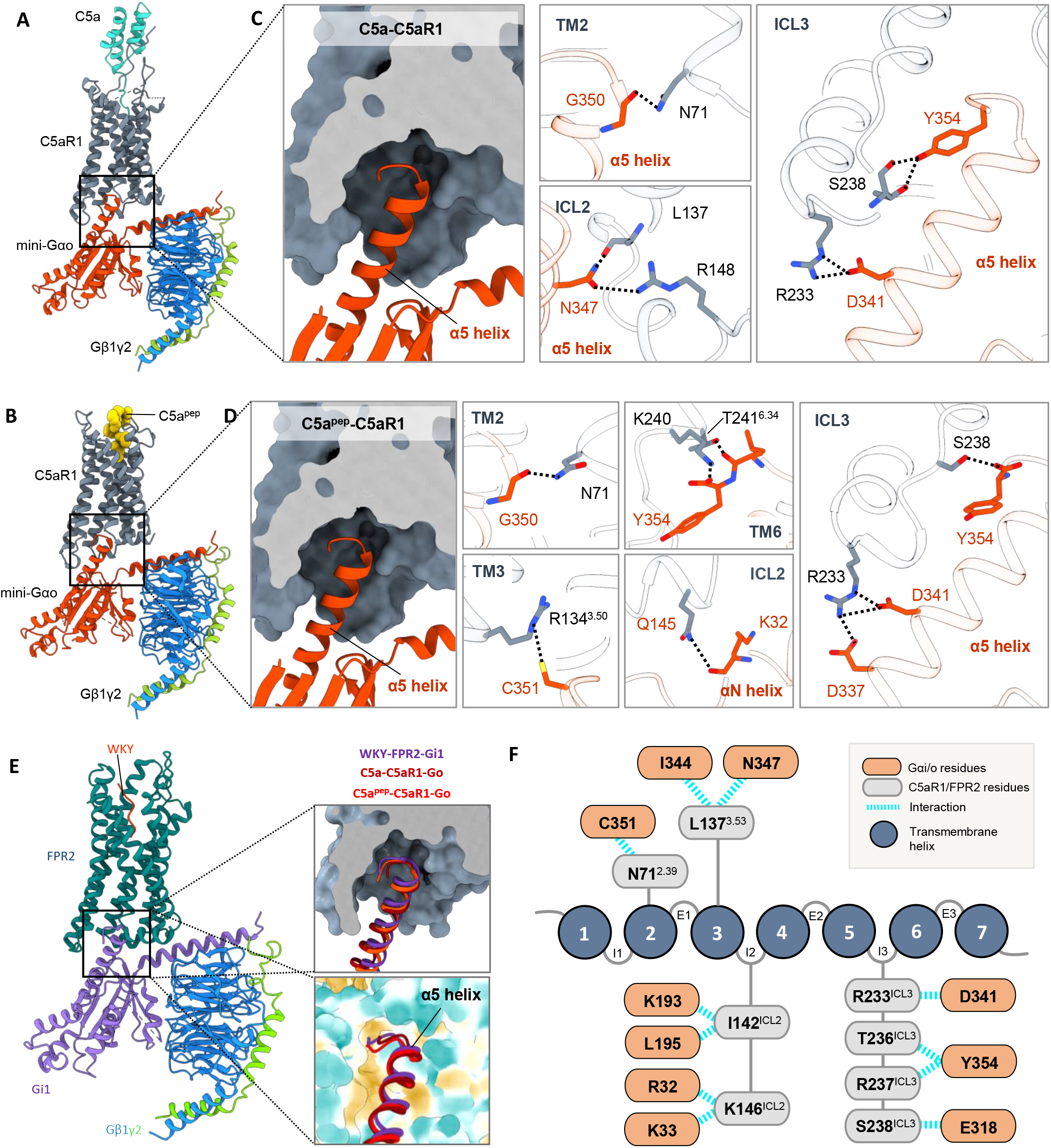
Overall interface of C5aR1-G-protein interaction. **(A, B)** Domain organization of heterotrimeric G-proteins in complex C5a/C5a^pep^-C5aR1 respectively. **(C)** The C-terminal α5 helix of Gαo docks into the cytoplasmic core of C5aR1 in the C5a-C5aR1-Go structure. Key interactions between residues of G-protein with residues of TMs, ICL2 and ICL3 of C5a bound C5aR1. **(D)** The C-terminal α5 helix of Gαo docks into the cytoplasmic core of C5aR1 in the C5a^pep^-C5aR1-Go structure. Key interactions between residues of G-protein with residues of TMs, ICL2 and ICL3 of C5a^pep^ bound C5aR1. **(E)** Comparative analysis of FPR2-Gi (PDB ID: 6OMM) with C5aR1-Go. The α5 helix of G-proteins inserts into a similar cavity (surface representation: top, hydrophobic surface representation: bottom) at the cytoplasmic face of the receptors. **(F)** Schematic representation of common residues of G-protein interacting with the residues of FPR2 and C5aR1. The respective residues mentioned are of C5aR1.

### Agonist-induced activation of C5aR1

In order to identify the conformational changes associated with C5aR1 activation, we compared these structures with previously determined crystal structures of C5aR1 in antagonist-bound state (Figure 9A)^31,37^. As expected, C5aR1 exhibits the major hallmarks of GPCR activation including a large outward movement of TM6 by about 8Å (as measured from Cα of Ser238^6.30^), and an inward movement of TM7 by about 6Å (as measured from Cα of Gly305^7.57^) (Figure 9B-C). In the inactive state structure of C5aR1, helix 8 exhibits an inverted orientation and it is sandwiched between TM1 and TM7, however, upon activation, it undergoes almost a 180º rotation (Figure 9B-C). Considering the G-protein interaction interface, this significant rotation and repositioning of helix 8 would be deemed essential to facilitate G-protein-coupling to the receptor. Taken together, these interactions promote the opening of a cavity towards the cytoplasmic face of the receptor, capable of accommodating the signal-transducers such as G-protein and the core interaction with βarrs (Figure 9D). In addition, a triad of conserved residues Ile^3.40^-Pro^5.50^-F^6.44^ was observed to form a hydrophobic interaction network which stabilized the inactive conformation of C5aR1 in the antagonist-bound state by preventing the movement of TM5 and TM6^31,37^ (Figure 9E-F). We observe rotameric shifts for these residues resulting in the opening of this hydrophobic triad and allowing the interaction with the α5 helix of Gαo (Figure 9E-F). Interestingly, C5aR1 harbors a slight variation of the highly conserved D^3.49^-R^3.50^-Y^3.51^ motif where Y^3.51^ is substituted with F^3.51^, and we observe a significant conformation change in this region upon receptor activation. In particular, the ionic interactions between D^3.49^ and R^3.50^, and R^3.50^ and S^6.31^ are disrupted upon receptor activation (Figure 9E-F). Finally, the other activation microswitches^38^ in C5aR1 such as the NPxxY, C(F)WxP and PIF motif also display noticeable conformational changes upon activation as observed in other GPCR-G-protein structures (Figure 9E-F). Although we have used C5a^pep^-C5aR1-Go structure for interpretation of activation-induced conformational changes considering the higher resolution; we note that these conformational changes are similar in the C5a-C5aR1-Go structure as well.

**Figure 9.**
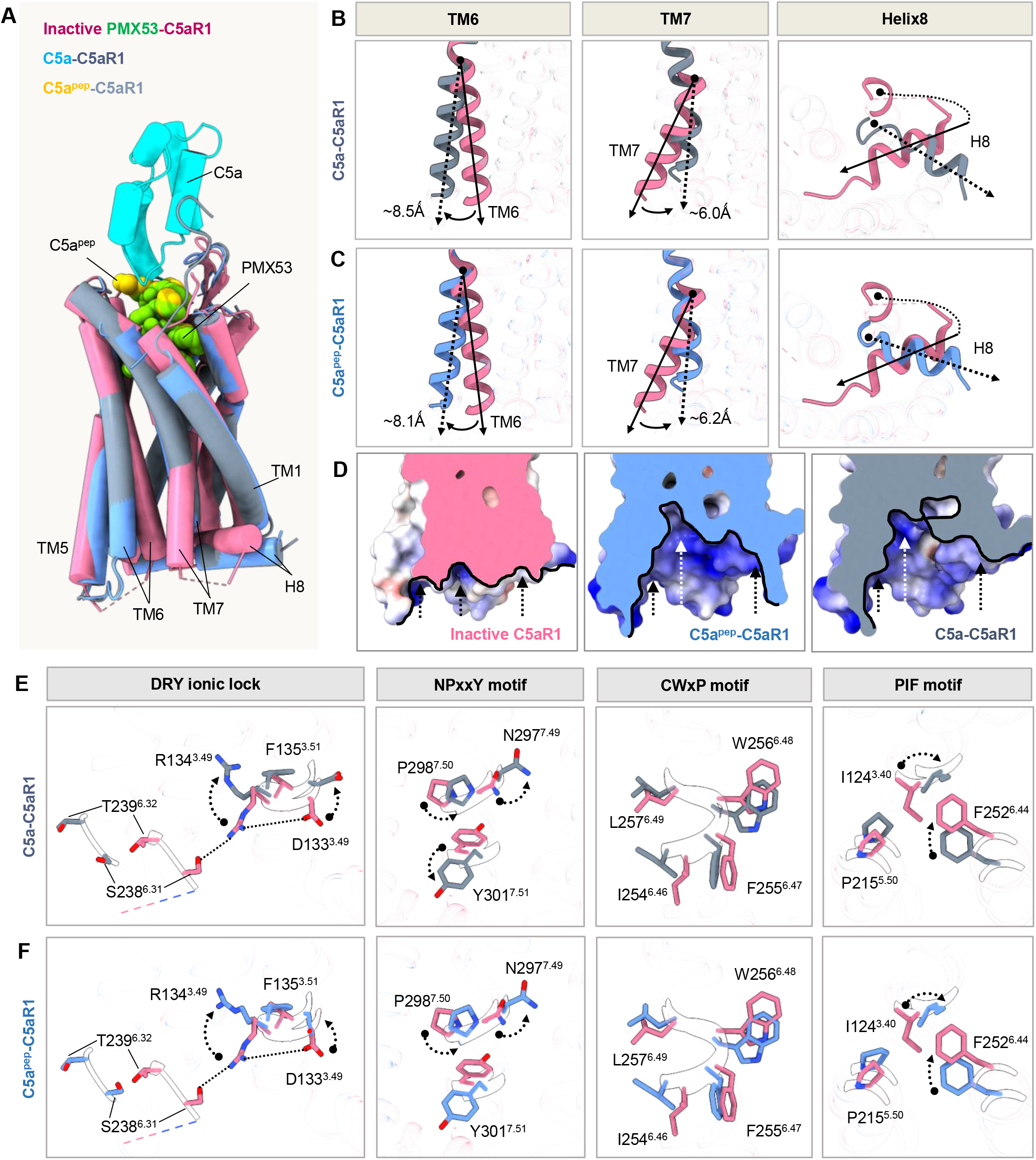
Activation-dependent conformational changes in C5aR1. **(A)** Structural alignment of the inactive (PMX53 bound C5aR1; PDB ID:6C1R) and active C5a and C5a^pep^ bound C5aR1. **(B, C)** Displacements of TM6, TM7 and helix 8 upon C5aR1 activation in C5a and C5a^pep^ bound C5aR1 structures respectively. **(D)** Opening of the cytoplasmic cavity in the active state structure of C5aR1. **(E, F)** Conformational changes in the conserved microswitches: (DRY(F), NPxxY, C(F)WxP(L), and PIF) upon C5aR1 activation.

### Concluding remarks

Our study provides the structural details of molecular interactions that are responsible for the conformational changes in C5aR1 upon activation, and the interface with G-proteins. These structural templates also provide a starting point for rational site-directed mutagenesis of the receptor to identify structural determinants of ligand bias, and potentially novel biased ligands as well. It is important to note that C5aR1 also exhibits moderate but significant secondary coupling to Gαq and Gα16 sub-types of G-proteins^9,39^, and additional structural snapshots in future studies may shed light on G-protein sub-type selectivity. Finally, structure determination of C5aR1 in complex with βarrs in subsequent studies should also facilitate a comprehensive understanding of receptor-transducer coupling and mechanisms that control receptor downregulation.

## Data availability statement

Any additional information required to reanalyze the data reported in this paper is available from the corresponding author upon reasonable request.

## Acknowledgements

This work is supported primarily by an extramural grant from the Department of Biotechnology (DBT) (BT/PR29041/BRB/10/1697/2018) sanctioned under the Membrane Protein Structural Biology initiative, and National Bioscience Award (BT/HRD/NBA/39/06/2018-19). In addition, the research in A.K.S.’s laboratory is supported by the Senior Fellowship of the DBT Wellcome Trust India Alliance (IA/S/20/1/504916) awarded to A.K.S., Science and Engineering Research Board (EMR/2017/003804, SPR/2020/000408, and IPA/2020/000405), Council of Scientific and Industrial Research [37(1730)/19/EMR-II], Indian Council of Medical research (F.NO.52/15/2020/BIO/BMS), Young Scientist Award from Lady Tata Memorial Trust, and IIT Kanpur. A.K.S. is EMBO Young Investigator. SS is supported by the Prime Minister’s Research Fellowship. We also thank Mithu Baidya for construct design of mC5aR1, Minakshi Baruah for initial purification ScFv16, and Ashutosh Ranjan for assistance in purification of C5a. Cryo-EM was performed at BioEM lab of the Biozentrum at the University of Basel, and we thank Carola Alampi and David Kalbermatter for their excellent technical assistance.

## Authors’ contribution

SS and MKY expressed and purified C5aR1, and reconstituted the complex with G-proteins for negative-staining and cryo-EM; SS carried out the functional assays with help from PS; JM and RB carried out negative-staining analysis and analyzed the cryo-EM data; VS, SM, SuM contributed in purification of ScFv16 and C5a; CS and SaS contributed in purification of G-proteins; MC collected cryo-EM data; JM prepared the figures together with SS, RB, MKY and MG; AKS supervised and managed the overall project; all authors contributed to data analysis, interpretation and manuscript writing.

## Accession number

The cryo-EM maps and structures have been deposited in the EMDB and PDB with accession numbers EMD-34943 and PDB ID: 8HPT (C5a^pep^-C5aR1-Go), and EMD-34947 and PDB ID: 8HQC (C5a-C5aR1-Go) respectively.

## Materials and Methods

### General reagents, plasmids, and cell culture

Most of the general reagents were purchased from Sigma Aldrich unless otherwise mentioned. Dulbecco′s Modified Eagle′s Medium (DMEM), Phosphate Buffer Saline (PBS), Trypsin-EDTA, Fetal-Bovine Serum (FBS), Hank’s Balanced Salt Solution (HBSS), and Penicillin-Streptomycin solution were purchased from Thermo Fisher Scientific. HEK293T cells (ATCC) were maintained in DMEM (Gibco, Cat. no: 12800-017) supplemented with 10% (v/v) FBS (Gibco, Cat. no: 10270-106) and 100U ml^-1^ penicillin and 100µg ml^-1^ streptomycin (Gibco, Cat. no: 15140122) at 37°C under 5% CO_2_. *Sf9* cells were obtained from Expression Systems and maintained in protein-free cell culture media purchased from Expression Systems (Cat. no: 96-001-01) at 27°C with 135 rpm shaking. The cDNA coding region of mC5aR1 was cloned in pcDNA3.1 vector with an N-terminal FLAG tag and in pVL1393 vector with an N-terminal FLAG tag followed by the N-terminal region of M4 receptor (residues 2-23). Constructs used for various NanoBiT assays were previously described^40^. All DNA constructs were verified by sequencing from Macrogen. Recombinant human C5a was purified from *E. coli* as previously described^11,12^. C5a^pep^ was synthesized from GenScript and its details are as previously described^11^. C5aR1 mutant was generated using NEB Q5 Site-Directed Mutagenesis Kit (NEB, Cat. no: E0554S).

### GloSensor-based cAMP assay

Ligand-induced Gαi-mediated inhibition of cAMP was measured using the GloSensor Assay, as previously described^22^. Briefly, HEK293T cells were co-transfected with FLAG-tagged receptor (3.5μg) and luciferase-based 22F cAMP biosensor construct (3.5μg) (Promega, Cat. no: E2301) using polyethyleneimine (PEI) (Polysciences, Cat. no: 19850) at a ratio of 3:1 (PEI:DNA 3:1). 14-16h after transfection, the cells were detached by trypsinization, resuspended in the assay buffer (20mM HEPES pH 7.4 and 1X HBSS) containing 0.5mg ml^-1^ of D-luciferin (GoldBio, Cat. no: LUCNA-1G) and seeded in 96-well white plates (SPL Life Sciences) at a density of 125,000 cells per well. The plates were then incubated for 1.5h at 37°C followed by an additional 30min at room temperature after which basal luminescence was measured. Since we were measuring Gαi activity, the cells were first treated with 5µM forskolin and luminescence was recorded using a microplate reader (FLUOStar Omega, BMG Labtech) till the readings stabilized (5-10 cycles) and then ligand was added at the indicated final concentration. Change in luminescence signal was recorded for 30 cycles. Data were normalized by treating luminescence observed at lowest concentration of agonist as 100%; plotting and analysis was performed using nonlinear regression in GraphPad Prism 9 software.

### NanoBiT-based βarr recruitment assay

Recruitment of βarrs to the receptor in response to agonist treatment was measured using the NanoBiT assay, as previously described. Briefly, HEK293T cells were co-transfected with either 2.5μg of C-terminally SmBiT-tagged mC5aR1 or 3.5μg of C-terminally SmBiT-tagged hC5aR1, and 3.5μg of βarr1/2 containing an LgBiT tag on their N-terminal end. 14-16h post-transfection, cells were harvested by trypsinization, resuspended in the assay buffer (5mM HEPES pH 7.4, 1X HBSS and 0.01% w/v BSA) containing 10µM of coelenterazine (GoldBio, Cat. no: CZ05) and seeded in 96-well white plates (SPL Life Sciences) at a density of 125,000 cells per well. After 1.5h of incubation at 37°C and 30min at room temperature, basal luminescence was recorded using a microplate reader (FLUOStar Omega, BMG Labtech). This was followed by addition of ligand at indicated final concentration and measurement of changes in signal for 20 cycles. Average data from 5 cycles was used for analysis. Data was normalized by calculating the fold increase in luminescence with respect to the signal observed at lowest concentration; plotting and analysis was performed using nonlinear regression in GraphPad Prism 9 software.

### Agonist-induced endosomal trafficking of βarrs

Agonist-induced endosomal trafficking of βarrs was measured as a surrogate for measuring receptor endocytosis. Briefly, HEK293T cells were co-transfected with 3µg of either mC5aR1 or hC5aR1, 2µg of βarr1/2 harboring an N-terminal SmBiT tag and 5µg of LgBiT-FYVE. For measuring βarr trafficking downstream of C5aR1^Tyr178Arg^ mutant, the following amounts of receptor DNA were used for transfection: 3µg of C5aR1^WT^ and 2µg of C5aR1^Tyr178Arg^. The rest of the protocol followed was same as described for measuring βarr recruitment, and data were normalized as mentioned above.

### Surface expression assay

Receptor surface expression was measured by performing whole-cell ELISA, as previously described^41^. Briefly, HEK293T cells transfected with the FLAG-tagged receptor of interest were seeded into 24-well plates (pre-coated with 0.01% poly-D-Lysine) at a density of 0.1 million cells per well, 24h after transfection. The next day, media was removed from the wells and the cells were washed once with ice-cold 1XTBS, followed by fixation with 4% (w/v) paraformaldehyde (prepared in 1XTBS) for 20min. The cells were then washed with 1XTBS extensively and blocking of non-specific sites was performed for 1.5h at room temperature by incubating with 1% BSA (w/v) prepared in 1XTBS. This was followed by incubation with anti-FLAG M2-HRP antibody (Sigma, Cat no. A8592) at a dilution of 1:5000 prepared in 1% BSA for another 1.5h. Excess unbound antibody was removed by washing the cells three times with 1% BSA. Cells were then incubated with 200μl of TMB (Thermo Scientific, Cat. no: 34028) substrate till a light blue color appeared and the reaction was stopped by pipetting 100μl of this colored solution to a 96-well plate already containing 100μl of 1M H_2_SO_4_. Absorbance was measured at 450nm in a multi-mode plate reader (Victor X4, Perkin Elmer). The remaining solution was removed from the wells and the cells were washed once with 1XTBS followed by incubation with 200μL of 0.2% (w/v) Janus green B (Sigma, Cat. no. 201677), a mitochondrial stain, for 20min. Cells were then washed extensively with deionized water to remove excess stain. 800μl of 0.5N HCl was added to each well to elute bound stain. 200μl of this solution was then transferred to a 96-well plate and absorbance was measured at 595nm. Data were normalized by calculating the ratio of A450 to A595.

### Expression and purification of C5a and C5aR1

Gene encoding C5a was cloned in pET-32a(+) vector with a Trx-6X-His tag at the N-terminal end and purified following previously described protocol with slight modification^11,12^. After Ni-NTA purification, we directly proceeded to TEV cleavage followed by cation-exchange chromatography. Codon-optimized mC5aR1 was expressed in *Spodoptera frugiperda* (*Sf9*) cells using baculovirus expression system with an N-terminal FLAG tag to facilitate purification. The receptor was purified as described previously^9^. Briefly, 72h post-infection, insect cells were harvested and lysed by sequentially douncing in low salt buffer (20mM HEPES pH 7.4, 10mM MgCl_2_, 20mM KCl, 1mM PMSF, and 2mM Benzamidine), high salt buffer (20mM HEPES pH 7.4, 1M NaCl, 10mM MgCl_2_, 20mM KCl, 1mM PMSF, and 2mM Benzamidine), and lysis buffer (20mM HEPES pH 7.4, 450mM NaCl, 2mM CaCl_2_, 1mM PMSF, 2mM Benzamidine and 2mM Iodoacetamide). After lysis, receptor was solubilized in 0.5% L-MNG (Anatrace, Cat. no: NG310) and 0.1% cholesteryl hemisuccinate (Sigma, Cat. no: C6512) for 2h at 4°C, under constant stirring. Post-solubilization, salt concentration was lowered to 150mM, and the receptor was purified on M1-FLAG column. After binding, FLAG beads were washed alternately with three washes of low salt buffer (20mM HEPES pH 7.4, 2mM CaCl_2_, 0.01% CHS, 0.01% L-MNG) and two washes of high salt buffer (20mM HEPES pH 7.4, 450mM NaCl, 2mM CaCl_2_, 0.01% L-MNG) to remove non-specific proteins. The bound receptor was eluted with FLAG elution buffer (20mM HEPES pH 7.4, 150mM NaCl, 0.01% MNG, 2mM EDTA, and 250µg ml^-1^ FLAG peptide) and alkylated with iodoacetamide to prevent aggregation. The purified receptor was concentrated using a 30kDa MWCO concentrator and stored at -80°C in 10% glycerol till further use. 100nM of hC5a or 1µM of C5a^pep^ were kept in all steps of receptor purification.

### Expression and purification of G-proteins

Gene for miniGαo1 subunit was cloned in pET-15b(+) vector with an in-frame 6X-His tag at the N-terminal end and expressed in *E. coli* BL21 (DE3) cells^42,43^. A starter culture supplemented with 0.2% glucose was grown in LB media at 37°C for 6-8h at 220 rpm, followed by overnight primary culturing at 30°C with 0.2% glucose supplementation. 15ml primary culture was inoculated in 1.5L TB (Terrific Broth) media and induced with 50μM IPTG at an O.D_600_ of 0.8 and cultured at 25°C for 18-20h. Cells were lysed in lysis buffer (40mM HEPES pH 7.4, 100mM NaCl, 10mM Imidazole, 10% Glycerol, 5mM MgCl_2_, 1mM PMSF, 2mM Benzamidine) in the presence of 1 mg ml^-1^ lysozyme, 50μM GDP and 100μM DTT. Cell debris was pelleted down by centrifuging at 18000 rpm for 30 mins at 4°C. Protein was enriched on Ni-NTA bead and after washing extensively with wash buffer (20mM HEPES pH 7.4, 500mM NaCl, 40mM Imidazole, 10% Glycerol, 50μM GDP and 1mM MgCl_2_), eluted with elution buffer (20mM HEPES pH 7.4, 100mM NaCl, 10% Glycerol, 500mM Imidazole). Eluted protein was pooled and stored at -80°C in 10% glycerol till further use.

The gene encoding the Gβ1 subunit with an in-frame C-terminal 6X-His tag and Gγ2 subunit was expressed in *Sf9* cells using the baculovirus expression system. Post 72h of infection, cells were harvested and resuspended in lysis buffer (20mM Tris-Cl pH 8.0, 150mM NaCl, 10% Glycerol, 1mM PMSF, 2mM Benzamidine, 1mM MgCl_2_). Cells were lysed by douncing and centrifuged at 18000 rpm for 40 mins at 4°C. Pellet was resuspended and dounced in solubilization buffer (20mM Tris-Cl pH 8.0, 150mM NaCl, 10% Glycerol, 1% DDM, 5mM β-ME, 10mM Imidazole, 1mM PMSF and 2mM Benzamidine) and solubilized at 4°C under constant stirring for 2h. Cell debris was pelleted down by centrifuging at 20000 rpm for 60min at 4°C. Protein was enriched on Ni-NTA resin, and after extensive washing with wash buffer (20mM Tris-Cl pH 8.0, 150mM NaCl, 30mM Imidazole, 0.02% DDM), the protein was eluted with elution buffer (20mM Tris-Cl pH 8.0, 150mM NaCl, 300mM Imidazole, 0.01% MNG). Eluted protein was concentrated with a 10kDa MWCO concentrator (Cytiva, Cat. no: GE28-9322-96) and stored at -80°C with 10% glycerol.

### Expression and purification of ScFv16

Gene encoding ScFv16^44^ was cloned in pET-42a(+) vector with an in-frame N-terminal 10X-His-MBP tag followed by a TEV cleavage site and expressed in *E. coli* Rosetta (DE3) strain^45^. Overnight primary culture was sub-cultured in 1L 2XYT media supplemented with 0.5% glucose and 5mM MgSO_4_. At O.D_600_ 0.6, culture was induced with 250µM IPTG for 16-18h at 18°C. Cells were resuspended in 20mM HEPES pH 7.4, 200mM NaCl, 10mM Imidazole, 2mM Benzamidine, 1mM PMSF and incubated at 4°C for 1h with constant stirring. Cells were disrupted by ultrasonication, and cell debris was removed by centrifugation at 18000 rpm for 40min at 4°C. Protein was enriched on Ni-NTA resins and nonspecifically bound proteins were removed by extensive washing (20mM HEPES pH 7.4, 200mM NaCl, 10mM Imidazole). Bound protein was eluted in elution buffer (20mM HEPES pH 7.4, 200mM NaCl, 300mM Imidazole). Subsequently, Ni-NTA elute was enriched on amylose resin (NEB, Cat. no: E8021L), and washed with buffer (20mM HEPES pH 7.4, 200mM NaCl) to remove nonspecific proteins. Protein was eluted with 10mM maltose (prepared in 20mM HEPES pH 7.4, 200mM NaCl), and the His-MBP tag was removed by overnight treatment with TEV protease. Tag-free ScFv16 was recovered by passing TEV-cleaved protein through Ni-NTA resin. Eluted protein was concentrated and cleaned by size exclusion chromatography on Hi-Load Superdex 200 preparative grade 16/600 column (Cytiva Life sciences, Cat. no: 17517501). Purified protein was flash frozen and stored at -80°C with 10% glycerol.

### Reconstitution of the C5a/C5a^pep^-C5aR1-Gαoβ1γ2-ScFv16 complexes

Purified mC5aR1 was incubated with 1.2 molar excess of Gαo1, Gβ1γ2, and ScFv16 at room temperature for 2h in the presence of 25mU ml^-1^ apyrase (NEB, Cat. no: M0398L) and either hC5a or C5a^pep^ for complex formation. The G-protein complex was separated from unbound components by loading on Superose 6 increase 10/300 GL SEC column and analyzed on SDS page. Complex fractions were pooled and concentrated to ∼10mg ml^-1^ using a 100MWCO concentrator (Cytiva, Cat. no: GE28-9323-19) and stored at -80°C until further use.

### Negative stain electron microscopy and data processing of C5a-C5aR1-Go and C5a^pep^-C5aR1-Go complexes

Negative staining of C5a-C5aR1-Go and C5a^pep^-C5aR1-Go samples were performed with uranyl formate stain to verify complex formation and homogeneity in accordance with a previously published protocol^9^. Complexes were diluted to 0.02 mg ml^-1^, immediately dispensed on glow discharged carbon/formvar coated 300 mesh Cu grids (PELCO, *Ted Pella*) and blotted off after incubation for 1min using a filter paper. Negative staining was done by touching the grid on a first drop of freshly prepared 0.75% (w/v) uranyl formate solution and blotted off using a filter paper. This was followed by incubating the grid on a second drop of stain for 30s and allowed to air dry before placing it on a TEM specimen grid holder. Imaging and data collection was performed at 30,000x magnification with a FEI Tecnai G2 12 Twin TEM (LaB6) operating at 120kV and equipped with a Gatan CCD camera (4k x 4k). Data processing of the collected micrographs was performed with Relion 3.1.2^46^. More than 10,000 particles were autopicked, extracted with a box size of 280 px and subjected to reference free 2D classification to obtain the 2D class averages.

### Cryo-EM sample preparation and data acquisition

3µl of the purified complexes of C5a^pep^-C5aR1-Go or C5a-C5aR1-Go were applied onto glow discharged Quantifoil holey carbon grids (Au, R2/1 M300) and vitrified using a Vitrobot Mark IV (Thermo Fisher Scientific, USA) operating at 10ºC and maintained at 90% humidity. Data collection was performed with a Titan Krios electron microscope (Thermofisher Scientific, USA) operating at 300kV equipped with Gatan Energy Filter. Movies were recorded in counting mode with a Gatan K2 Summit direct electron detector DED (Gatan, USA) using the automated SerialEM software at a nominal magnification of 165 000x and a pixel size of 0.82Å at the specimen level. 24,711 movie stacks for C5a^pep^-C5aR1-Go and 22,014 movie stacks for C5a-C5aR1-Go consisting of 40 frames were collected with a defocus value in the range of 0.5 to 2.5µm with a total accumulated dose of 42 e^-^/A^2^ and total exposure time of 4s.

### Cryo-EM data processing

The flowchart for processing the vitrified C5a^pep^-C5aR1-Go and C5a-C5aR1-Go complexes are shown in Figures S3 and S4. All data processing steps were performed with cryoSPARC version 3.3.2 or version 4^24^. Briefly, 24,711 movies of C5a^pep^-C5aR1-Go were imported and subjected to Patch motion correction (multi) followed by CTF estimation with Patch CTF estimation (multi). 23,723 motion corrected micrographs with CTF fit resolution better than 6Å were selected for further processing. 1,886,363 particles were autopicked with the blob-picker sub-program within the cryoSPARC suite, extracted with a box size of 360 px and fourier cropped to 64 px (pixel size of 4.61) for reference free 2D classification. Several rounds of iterative 2D classification yielded class averages representing different orientations of the complex. A subset of 835,654 clean particles from the 2D classification were re-extracted with a box size of 360 px and fourier cropped to 256 px (pixel size of 1.15). This was followed by Ab-initio reconstruction and heterogenous refinement with C1 symmetry yielding 3 models. 380,463 particles corresponding to the class with clear complex conformation were re-extracted with full box size of 416 px, fourier cropped to 360 px and subjected to non-uniform refinement followed by local refinement with mask on the complex excluding the micelle. This led to a reconstruction at 3.45Å (voxel size of 0.9476) as determined by gold standard Fourier Shell Correlation (FSC) using the 0.143 criterion. Blocres sub-program within cryoSPARC version 3.3.2 was used to estimate local resolution of all reconstructions.

For the C5a-C5aR1-Go dataset, 22,014 movies were imported and subjected to Patch motion correction (multi). CTF estimation was performed on the motion corrected micrographs and 21,449 micrographs with CTF fit resolution better than 6Å were selected for downstream processing. Automated particle picking with blob-picker resulted in 2,601,754 particles which were extracted with a box size of 360 px and fourier cropped to box size of 64 (pixel size of 4.61). These particles were then subjected to several rounds of 2D classification and class averages with clear conformations of the complex were selected and extracted with a box size of 360px and fourier cropped to 256 px (pixel size of 1.15). These clean set of particles were subjected to Ab-initio reconstruction and heterogeneous refinement yielding 3 models. 173,416 particles corresponding to the 3D class with evident secondary features were re-extracted with full box size of 416px and fourier cropped to 360px. This was followed by non-uniform refinement and local refinement with mask on the complex resulting in a final map at 3.89Å resolution (voxel size of 0.9476) according to the gold standard Fourier shell correlation (FSC) criterion of 0.143. All maps were sharpened with “Autosharpen” sub-program within the Phenix suite^47^ for better visualization and model building.

### Model building and refinement

The receptor coordinates from the cryo-EM structure of human formyl peptide receptor 2 (PDB ID: 7WVV) and the coordinates for the Gαo, Gβ1, Gɣ2 from the cryo-EM structure of Muscarinic acetylcholine receptor 2-Go complex (PDB ID: 6OIK) were used as an initial model to dock into the EM density of C5a^pep^-C5aR1-Go complex using Chimera^48^. This was followed by manual rebuilding of the model along with the ligand in COOT^49^ and iterative real space refinement in Phenix^47^. This yielded a model with 95.13% in the most favoured region and 4.87% in the allowed region of the Ramachandran plot.

For the C5a-C5aR1-Go complex map, the coordinates of C5a^pep^-C5aR1-Go complex was used as an initial model and docked into the EM density with the “Fit in map” extension in Chimera. Similarly, the coordinates corresponding to human C5a were taken from a previously solved crystal structure of the human C5a in complex with MEDI7814, a neutralising antibody (PDB ID: 4UU9). The model so obtained was docked in Chimera, manually rebuilt in COOT and subjected to several rounds of real space refinement in Phenix to reach a final model with 95.87% in the favoured region and 4.13% in the allowed region of the Ramachandran plot. Data collection, 3D reconstruction and refinement statistics have been included as Figure S5. All figures were prepared either with Chimera or ChimeraX software^48,50^. Buried surface and interface surface area have been calculated with PDBePISA webserver^51^. Ligand-receptor interactions presented in Figure S12 were identified using PDBsum^52^.

