## Supplemental Material for "Structural insights into ligand-recognition, activation, and signaling-bias at the complement C5a receptor, C5aR1"

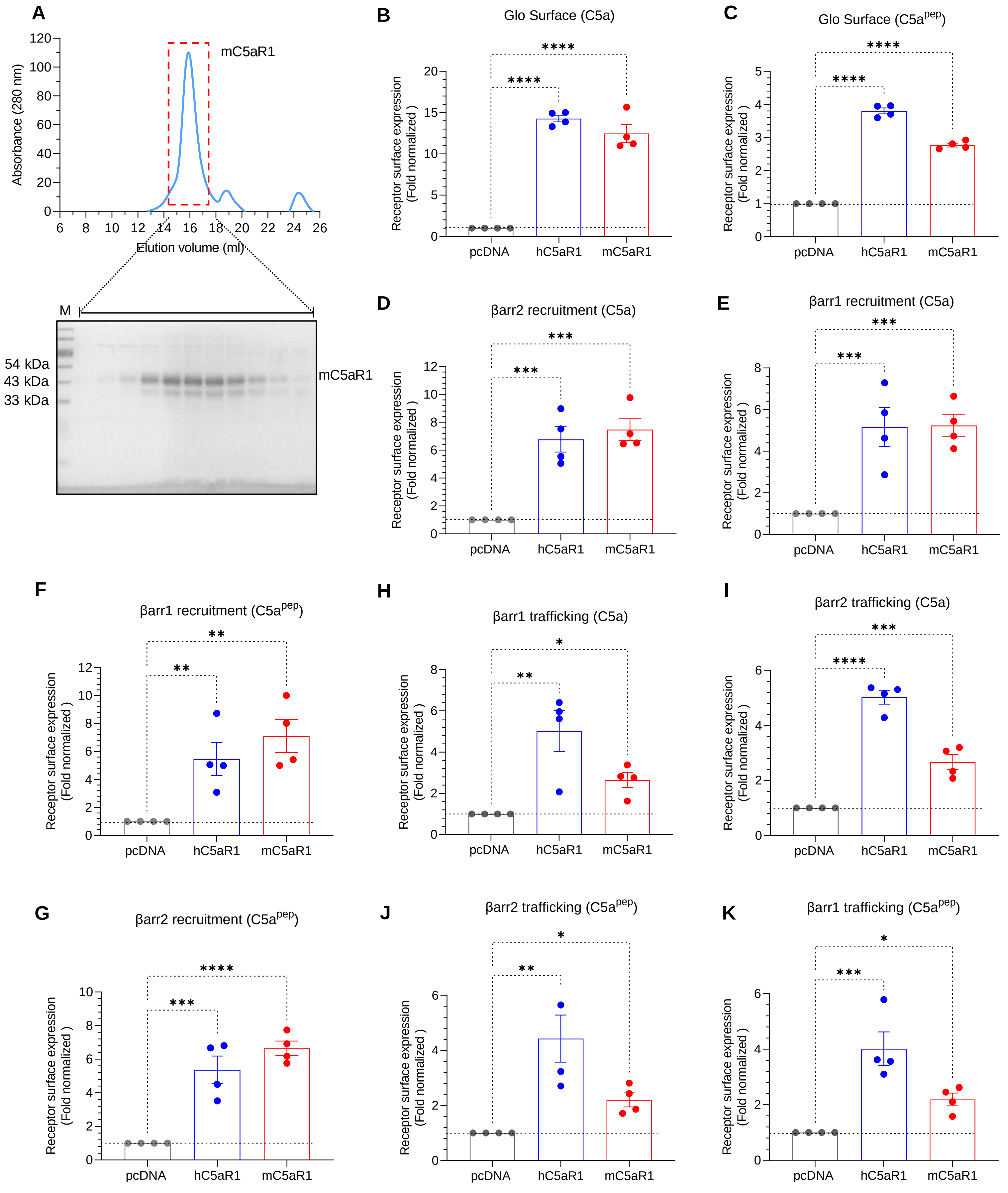
A
mC5aR1
Absorbance (280 nm)
Elution volume (ml)
M
54 kDa
43 kDa
33 kDa
mC5aR1
B
Glo Surface (C5a)
C
Glo Surface (C5a^pep)
D
βarr2 recruitment (C5a)
E
βarr1 recruitment (C5a)
F
βarr1 recruitment (C5a^pep)
G
βarr2 recruitment (C5a^pep)
H
βarr1 trafficking (C5a)
I
βarr2 trafficking (C5a)
J
βarr2 trafficking (C5a^pep)
K
βarr1 trafficking (C5a^pep)
Receptor surface expression (Fold normalized )
pcDNA
hC5aR1
mC5aR1

**Figure S1. Purification and surface expression profiles of C5aR1.**

**(A)** Size exclusion chromatography (top) and SDS-PAGE of mouse C5aR1 shows a homogeneous population following purification. **(B-K)** Surface expression of indicated receptors measured by using whole cell ELISA (mean±SEM; n=4; One-way ANOVA, Two-stage step-up method of Benjamini, Krieger and Yekutieli, *p<0.05, **p<0.01, ***p<0.001, ****p<0.0001) in various assays.

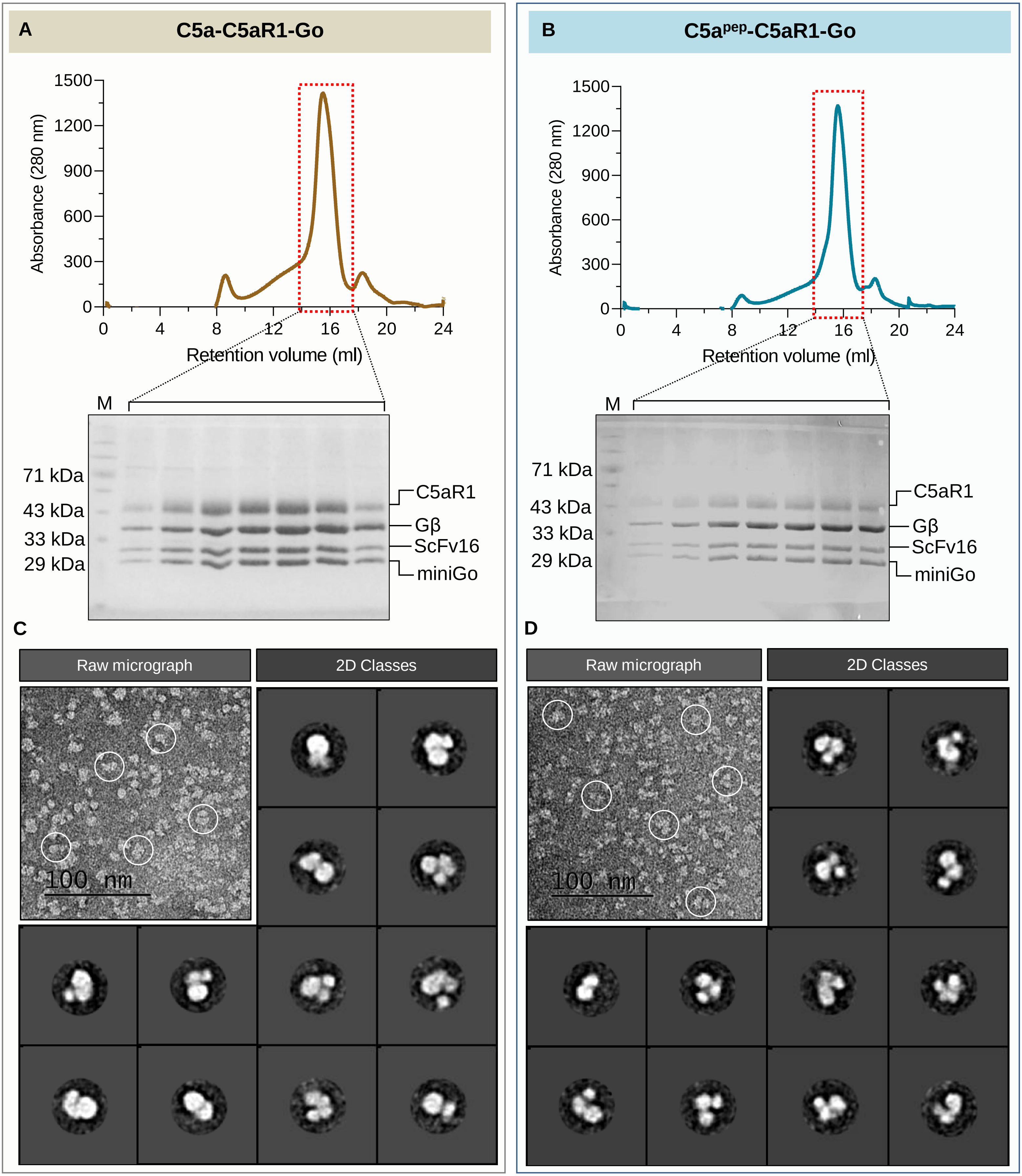
A
C5a-C5aR1-Go
Absorbance (280 nm)
1500
1200
900
600
300
0
0
4
8
12
16
20
24
Retention volume (ml)
M
71 kDa
43 kDa
33 kDa
29 kDa
C5aR1
Gβ
ScFv16
miniGo
B
C5a$^{pep}$-C5aR1-Go
Absorbance (280 nm)
Retention volume (ml)
M
71 kDa
43 kDa
33 kDa
29 kDa
C5aR1
Gβ
ScFv16
miniGo
C
Raw micrograph
2D Classes
100 nm
D
Raw micrograph
2D Classes
100 nm

**Figure S2. Purification of C5a/C5a$^{pep}$-C5aR1-Go complex and visualization through negative staining EM.**

**(A, B)** SEC chromatogram and SDS analysis of C5a-C5aR1-G-protein and C5a$^{pep}$-C5aR1-G-protein complex, respectively. SDS-PAGE shows relative abundance of the components following complex formation. **(C, D)** Negative staining raw micrographs and 2D class averages of C5a-C5aR1-G-protein and C5a$^{pep}$-C5aR1-G-protein complex, respectively.

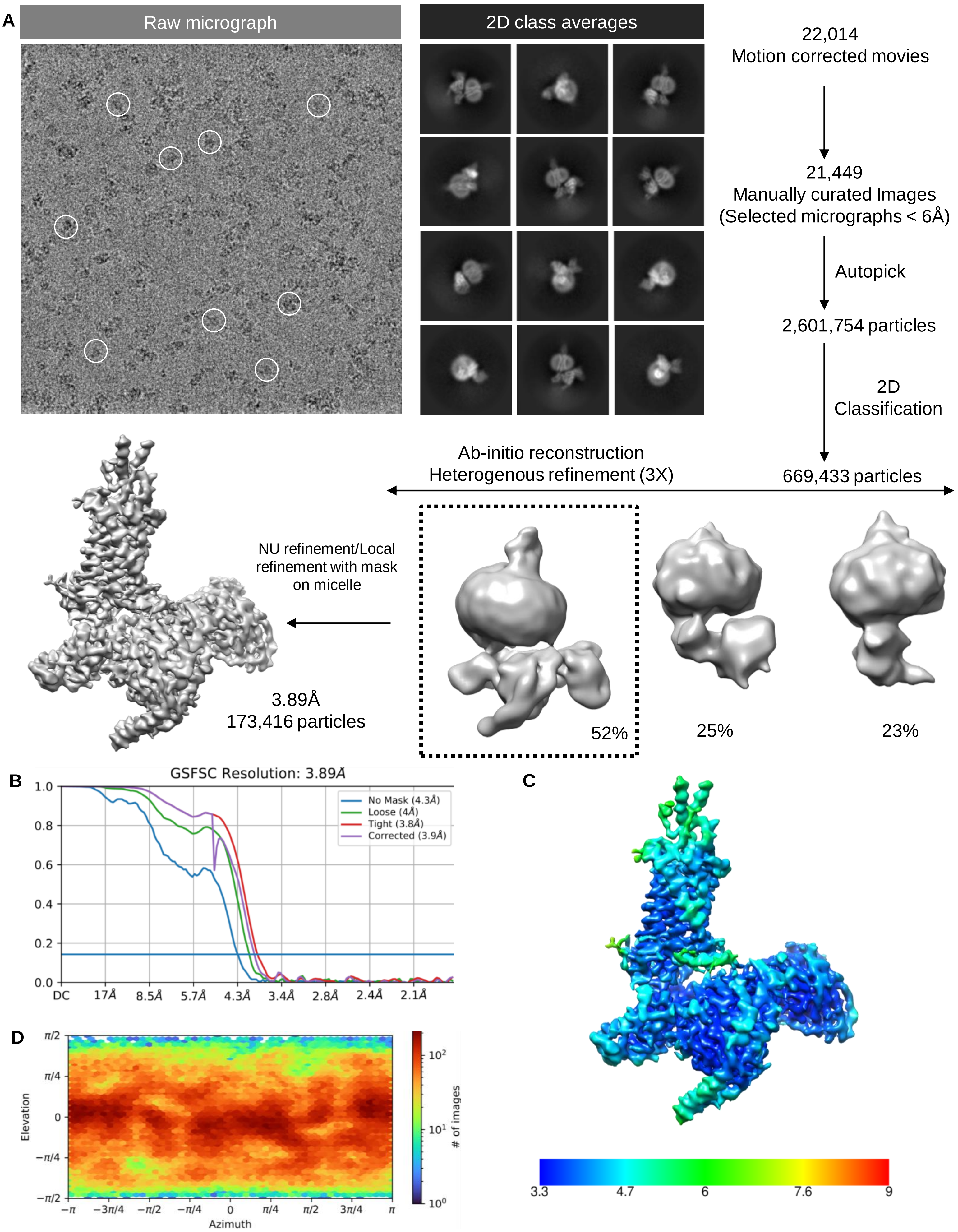
A
Raw micrograph
2D class averages
22,014
Motion corrected movies
21,449
Manually curated Images
(Selected micrographs < 6Å)
Autopick
2,601,754 particles
2D
Classification
669,433 particles
Ab-initio reconstruction
Heterogenous refinement (3X)
NU refinement/Local
refinement with mask
on micelle
3.89Å
173,416 particles
52%
25%
23%
B
GSFSC Resolution: 3.89Å
No Mask (4.3Å)
Loose (4Å)
Tight (3.8Å)
Corrected (3.9Å)
DC
17Å
8.5Å
5.7Å
4.3Å
3.4Å
2.8Å
2.4Å
2.1Å
C
3.3
4.7
6
7.6
9
D
Elevation
Azimuth
### of images

**Figure S3: Workflow for cryo-EM data processing of C5a-C5aR1-Go complex.**

**(A)** Schematic representation of cryo-EM data processing workflow. **(B)** Gold standard fourier shell correlation curve (GSFSC) at 0.143 threshold indicates an overall resolution of 3.89Å. **(C)** Local resolution map of the 3D reconstruction in front view. **(D)** Angular distribution of the particles used for final reconstruction of C5a-C5aR1-Go complex.

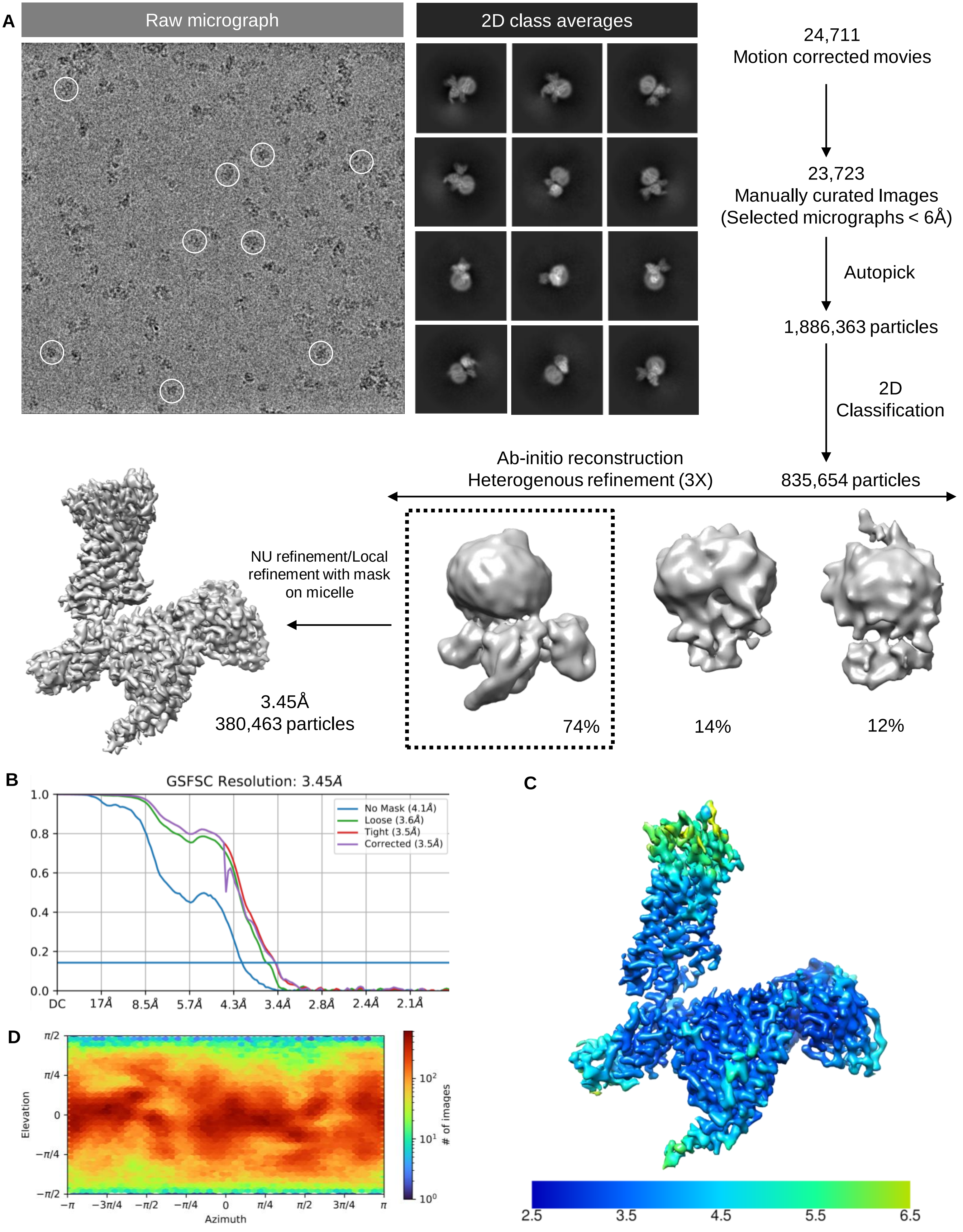
A
Raw micrograph
2D class averages
24,711
Motion corrected movies
23,723
Manually curated Images
(Selected micrographs < 6Å)
Autopick
1,886,363 particles
2D
Classification
Ab-initio reconstruction
Heterogenous refinement (3X)
835,654 particles
NU refinement/Local
refinement with mask
on micelle
3.45Å
380,463 particles
74%
14%
12%
B
GSFSC Resolution: 3.45Å
No Mask (4.1Å)
Loose (3.6Å)
Tight (3.5Å)
Corrected (3.5Å)
DC 17Å 8.5Å 5.7Å 4.3Å 3.4Å 2.8Å 2.4Å 2.1Å
C
2.5 3.5 4.5 5.5 6.5
D
Elevation
Azimuth
### of images

**Figure S4: Workflow for cryo-EM data processing of C5a$^{pep}$-C5aR1-Go complex.**

**(A)** Schematic representation of cryo-EM data processing workflow. **(B)** Gold standard fourier shell correlation curve (GSFSC) at 0.143 threshold indicates an overall resolution of 3.45Å. **(C)** Local resolution map of the 3D reconstruction in front view. **(D)** Angular distribution of the particles used for final reconstruction of C5a$^{pep}$-C5aR1-Go complex.

| Data collection and processing | | |
|---|---|---|
| | C5a$^{pep}$-C5aR1-Go PDB 8HPT, EMD-34943 | C5a-C5aR1-Go PDB 8HQC, EMD-34947 |
| Microscope | Titan Krios | Titan Krios |
| Camera | GIF/K2 | GIF/K2 |
| Magnification | 165,000x | 165,000x |
| Voltage (kV) | 300 | 300 |
| Defocus range (µm) | 0.5-2.5 | 0.5-2.5 |
| Exposure time (s) | 4 | 4 |
| Total dose ($e^-/A^2$) | 42 | 42 |
| Number of frames | 40 | 40 |
| Pixel size (Å) | 0.82 | 0.82 |
| Micrographs (no.) | 24,711 | 22,014 |
| Initial particles (no.) | 1,886,363 | 2,601,754 |
| Symmetry imposed | C1 | C1 |
| Final particles (no.) | 380,463 | 173,416 |
| Map resolution (Å) | 3.45 | 3.89 |
| FSC threshold | 0.143 | 0.143 |
| **Refinement** | | |
| Initial model (PDB Code) | 7WVV, 6OIK | 8HPT, 4UU9 |
| Model resolution (Å) | 3.7 | 4.1 |
| FSC threshold | 0.5 | 0.5 |
| **Model composition** | | |
| Non-hydrogen atoms | 7,893 | 8,538 |
| Protein residues | 1,094 | 1,186 |
| Ligand atoms | DAR=1 | 0 |
| **R. M.S. deviations** | | |
| Bond length (Å) | 0.006 | 0.003 |
| Bond angle (º) | 1.114 | 0.686 |
| **Validation** | | |
| Favored (%) | 95.13 | 95.87 |
| Allowed (%) | 4.87 | 4.13 |
| Disallowed (%) | 0 | 0 |
| MolProbity score | 1.59 | 1.65 |
| Clash Score | 4.81 | 6.6 |

**Figure S5:** 3D reconstruction and model refinement statistics.

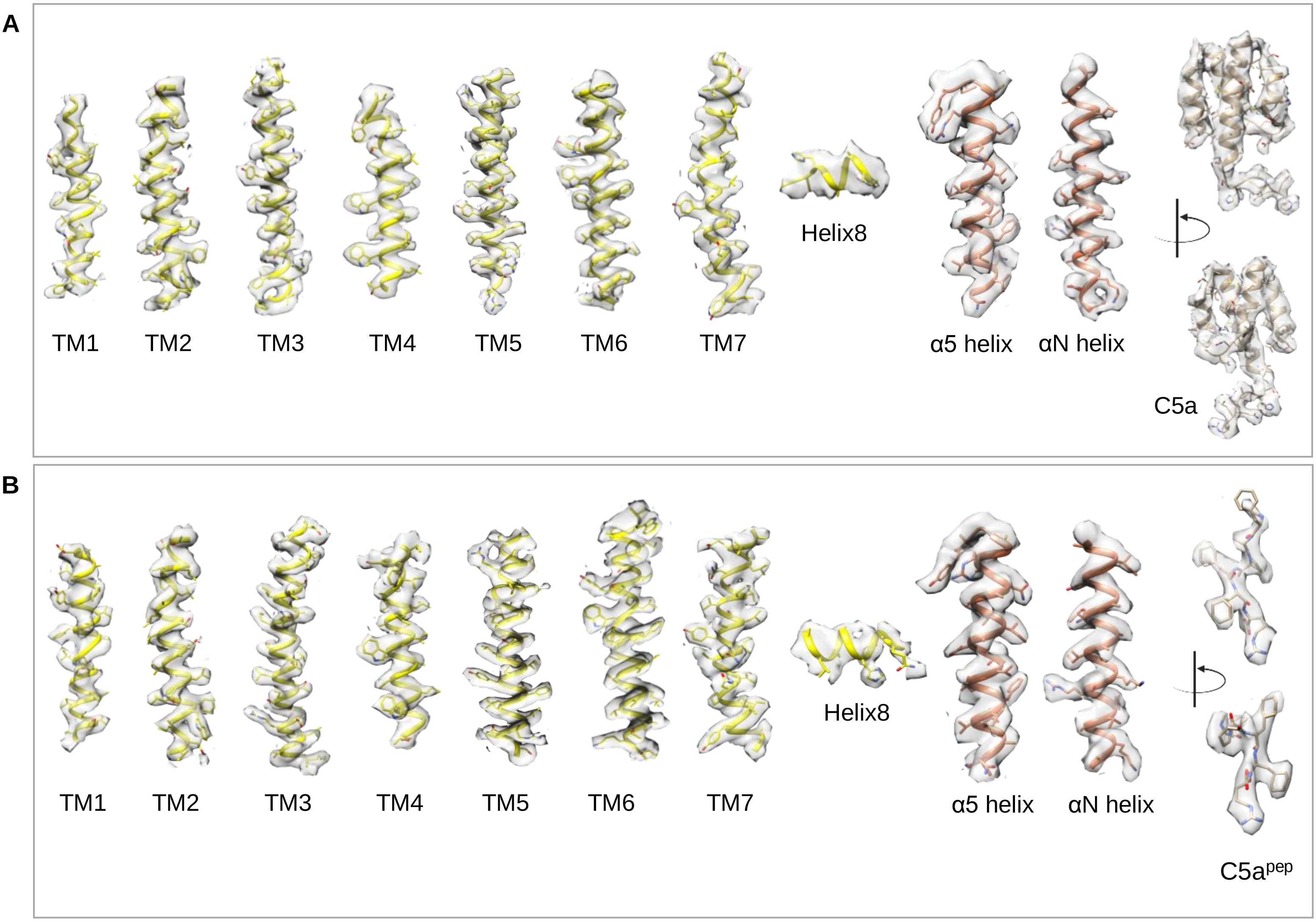
A
TM1
TM2
TM3
TM4
TM5
TM6
TM7
Helix8
α5 helix
αN helix
C5a
B
TM1
TM2
TM3
TM4
TM5
TM6
TM7
Helix8
α5 helix
αN helix
C5apep

**Figure S6: Representative electron density maps.**

**(A)** EM densities for the TMs of C5a-C5aR1-Go structure: TM1 to TM7, Helix8, α5 helix, αN helix, and C5a (left to right). **(B)** EM densities for the TMs of $C5a^{pep}$-C5aR1-Go structure: TM1 to TM7, Helix8, α5 helix, αN helix, and $C5a^{pep}$ (left to right).

| Component | Total residues | Resolved residues | Total residues | Resolved residues |
| --- | --- | --- | --- | --- |
| | C5a-C5aR1-Go | | $C5a^{pep}$-C5aR1-Go | |
| C5a/$C5a^{pep}$ | 1T-R74 | T1-R74 | 1NME-F-dR6 | 1NME-F-dR6 |
| C5aR1 | M1-V351 | $P24^{N\text{-}term}$-$W102^{ECL1}$<br>$A106^{3.22}$-$T186^{ECL2}$<br>$K201^{5.37}$-$S315^{H8}$ | M1-V351 | $G36^{1.31}$-$E65^{1.60}$<br>$A69^{2.37}$-$Y178^{ECL2}$<br>$K201^{5.37}$-$S315^{H8}$ |
| miniGαo | M1-H57<br>T172-Y366 | L5-I55<br>T182-D232<br>R243-Y354 | M1-H57<br>T172-Y366 | L5-K54<br>T183-Y231<br>M244-C325<br>D328-Y354 |
| Gβ | M1-N340 | E3-N340 | M1-N340 | E3-N340 |
| Gγ | M1-L71 | A7-R62 | M1-L71 | A7-R62 |
| ScFv16 | D1-K248 | D1-G122<br>S136-Y235<br>T238-L247 | D1-K248 | D1-S121<br>S136-K248 |

**Figure S7:** List of resolved residues in the cryo-EM structures of C5a and C5a$^{pep}$ bound C5aR1-Go compared to the total residues in the protein sequence used for structure analysis.

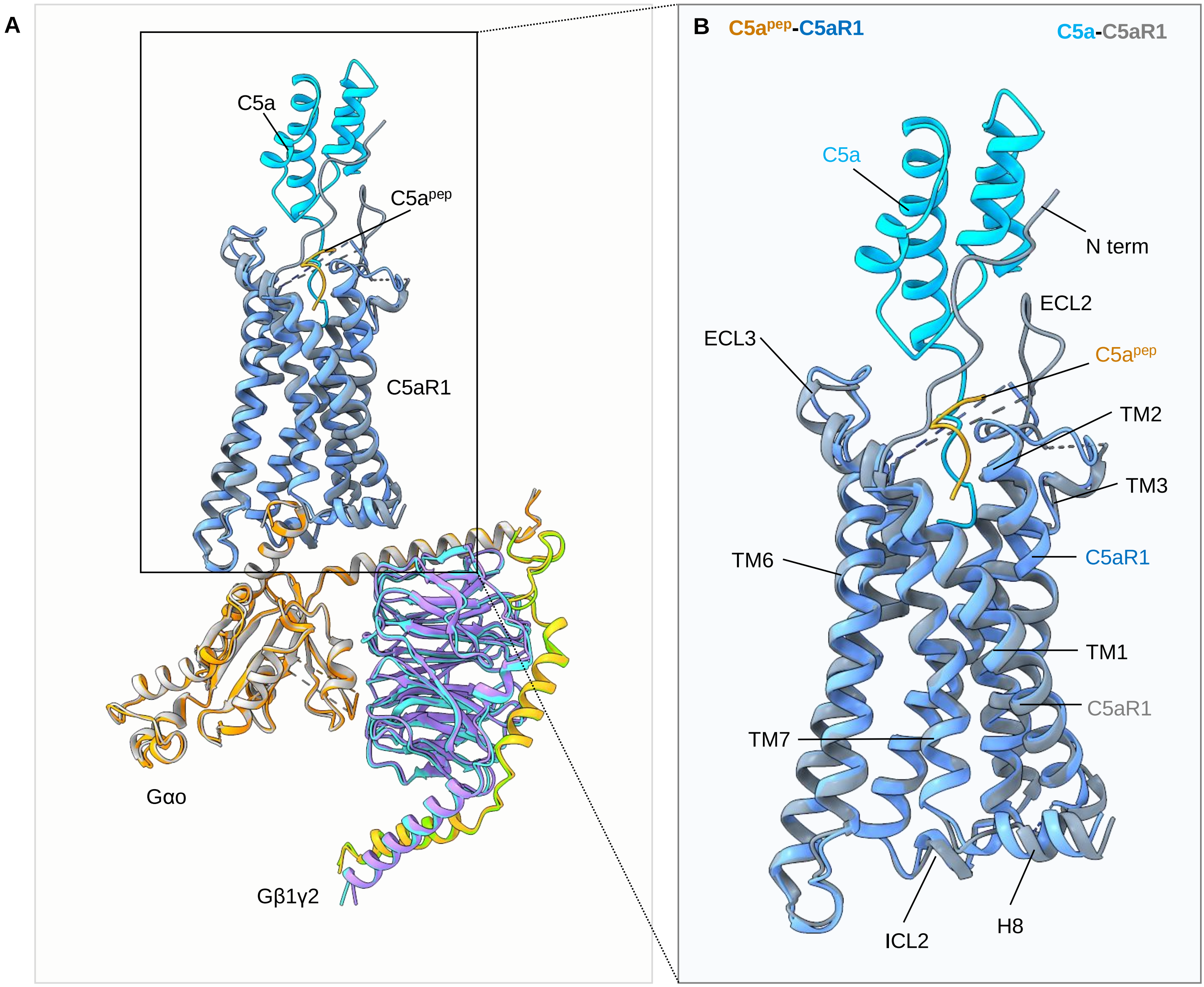
A
C5a
C5a^pep
C5aR1
Gαo
Gβ1γ2
B
C5a^pep-C5aR1
C5a-C5aR1
C5a
N term
ECL2
ECL3
C5a^pep
TM2
TM3
TM6
C5aR1
TM1
C5aR1
TM7
ICL2
H8

**Figure S8: Structural alignment of C5a/C5a$^{pep}$-C5aR1-Go complexes.**

**(A, B)** The overall structures of C5a and C5a$^{pep}$ bound C5aR1-Go complexes show similar architecture upon superposition of the whole complex and also of the ligand bound receptors, respectively.

**A**

| C5a-C5aR1 interface | | |
|---|---|---|
| **Chain A (C5aR1)** | **Distance (Å)** | **Chain D (C5a)** |
| Ile28 (N-terminal loop) | Arg40 (3.29), Ile41 (3.70) | Arg40, Ile41 |
| His29 (N-terminal loop) | Arg37 (2.83), Arg40 (3.44) | Arg37, Arg40 |
| Leu92 (TM2) | Gly73 (3.66) | Gly73 |
| Thr95 (TM2) | Leu72 (3.76) | Leu72 |
| Asn100 (ECL1) | Leu72 (3.74) | Leu72 |
| Pro113 (TM3) | Gln71 (3.47) | Gln71 |
| Met120 (TM3) | Gly73 (3.86) | Gly73 |
| Arg175 (ECL2) | Gln71 (3.71) | Gln71 |
| Glu176 (ECL2) | Ser66 (2.31), His67 (2.43) | Ser66, His67 |
| Phe181 (ECL2) | Ala26 (3.90), Cys27 (3.78) | Ala26, Cys27 |
| Tyr259 (TM6) | Arg74 (3.28) | Arg74 |
| Gly263 (TM6) | Arg74 (3.72) | Arg74 |
| Ile266 (TM6) | Arg74 (3.75) | Arg74 |
| Val287 (TM7) | Gly73 (3.81), Arg74 (3.87) | Gly73, Arg74 |

**B**

| $C5a^{pep}$-C5aR1 interface | | |
|---|---|---|
| **Chain A (C5aR1)** | **Distance (Å)** | **Chain D ($C5a^{pep}$)** |
| Leu92 (TM2) | DAR406 (3.68) | DAR406 |
| Ile116 (TM3) | DAR406 (3.42) | DAR406 |
| Leu117 (TM3) | DAR406 (3.23) | DAR406 |
| Arg175 (ECL2) | ALC405 (3.51) | ALC405 |
| Glu176 (ECL2) | MEA401 (3.62) | MEA401 |
| Tyr178 (ECL2) | MEA401 (3.07), ZAL404 (3.23), ALC405 (3.16) | MEA401, ZAL404, ALC405 |
| Glu280 (TM7) | Pro403 (3.54) | Pro403 |
| Asn283 (TM7) | DAR406 (3.28) | DAR406 |
| Val287 (TM7) | DAR406 (3.46) | DAR406 |

**Figure S9: (A, B)** List of C5a/C5a$^{pep}$-C5aR1 interactions.

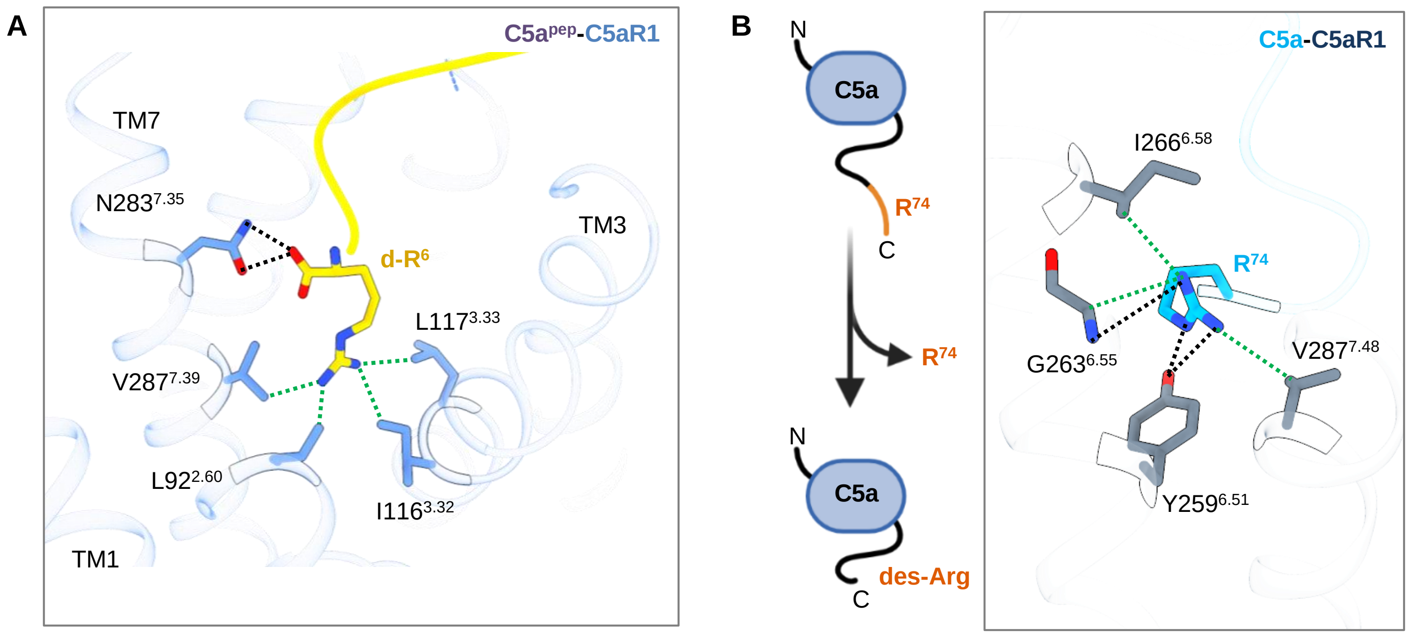
A
C5apep-C5aR1
TM7
N283[7.35]
d-R[6]
TM3
L117[3.33]
V287[7.39]
L92[2.60]
I116[3.32]
TM1
B
N
C5a
R[74]
C
R[74]
N
C5a
des-Arg
C
C5a-C5aR1
I266[6.58]
R[74]
G263[6.55]
V287[7.48]
Y259[6.51]

**Figure S10: Critical interactions of the terminal Arg of C5a/C5a$^{pep}$ with C5aR1.**

**(A)** D-Arg$^{6}$ makes extensive interactions with the residues of the extracellular core of the receptor in the C5a$^{pep}$-C5aR1 structure. **(B)** Proteolytic cleavage of the terminal Arg from C5a results in a natural variant, namely C5a$^{des\text{-}Arg}$. Terminal Arg$^{74}$ engages with various residues from the TMs of C5aR1. (Polar interactions: Black dotted lines, Non-bonded contacts: Green dotted lines).

```
            .........10.........20.........30.........40.........50
mC5aR1      MDPIDNSSFE I-NYDHYGTM DPNIPADGIH LPKRQPGDVA ALIIYSVVFL
hC5aR1      MDSFNYTTPD YGHYDDKDTL DLNTPVDKTS NTLRVP-DIL ALVIFAVVFL
Consistency **34515506 304**223*7 *1*3*5*233 132*2*0*83 **8*66****

            .........60.........70.........80.........90.........100
mC5aR1      VGVPGNALVV WVTAFEARRA VNAIWFLNLA VADLLSCLAL PVLFTTVLNH
hC5aR1      VGVLGNALVV WVTAFEAKRT INAIWFLNLA VADFLSCLAL PILFTSIVQH
Consistency ***1****** *******6*4 8********* ***4****** *8***5864*

            .........110.........120.........130.........140.........150
mC5aR1      NYWYFDATAC IVLPSLILLN MYASILLLAT ISADRFLLVF KPIWCQKVRG
hC5aR1      HHWPFGGAAC SILPSLILLN MYASILLLAT ISADRFLLVF KPIWCQNFRG
Consistency 45*0*344** 28******** ********** ********** ******43**

            .........160.........170.........180.........190.........200
mC5aR1      TGLAWMACGV AWVLALLLTI PSFVYREAYK DFYSEHTVCG INYGGGSFPK
hC5aR1      AGLAWIACAV AWGLALLLTI PSFLYRVVRE EYFPPKVLCG VDYSHDK-RR
Consistency 4****5**4* **1******* ***6**2525 66633246** 85*4134026

            .........210.........220.........230.........240.........250
mC5aR1      EKAVAILRLM VGFVLPLLTL NICYTFLLLR TWSRKATRST KTLKVVMAVV
hC5aR1      ERAVAIVRLV LGFLWPLLTL TICYTFILLR TWSRRATRST KTLKVVVAVV
Consistency *6****6**5 6**61***** 4*****7*** ****6***** ******5***

            .........260.........270.........280.........290.........300
mC5aR1      ICFFIFWLPY QVTGVMIAWL PPSSPTLKRV EKLNSLCVSL AYINCCVNPI
hC5aR1      ASFFIFWLPY QVTGIMMSFL EPSSPTFLLL KKLDSLCVSF AYINCCINPI
Consistency 32******** ****8*564* 3*****4226 5**5*****4 ******8***

            .........310.........320.........330.........340.........350
mC5aR1      IYVMAGQGFH GRLLRSLPSI IRNALSEDSV GRDSKTFTPS TTDTSTRKSQ
hC5aR1      IYVVAGQGFQ GRLRKSLPSL LRNVLTEESV VRESKSFTRS TVDTMAQKTQ
Consistency ***5*****3 ***26****7 7**5*5*6** 1*6**5**2* *4**345*5*

            ..
mC5aR1      AV
hC5aR1      AV
Consistency **
```

Unconserved 0 1 2 3 4 5 6 7 8 9 10 Conserved

**Figure S11:** Sequence alignment of human C5aR1 and mouse C5aR1 highlighting conservation and differences between the two sequences.

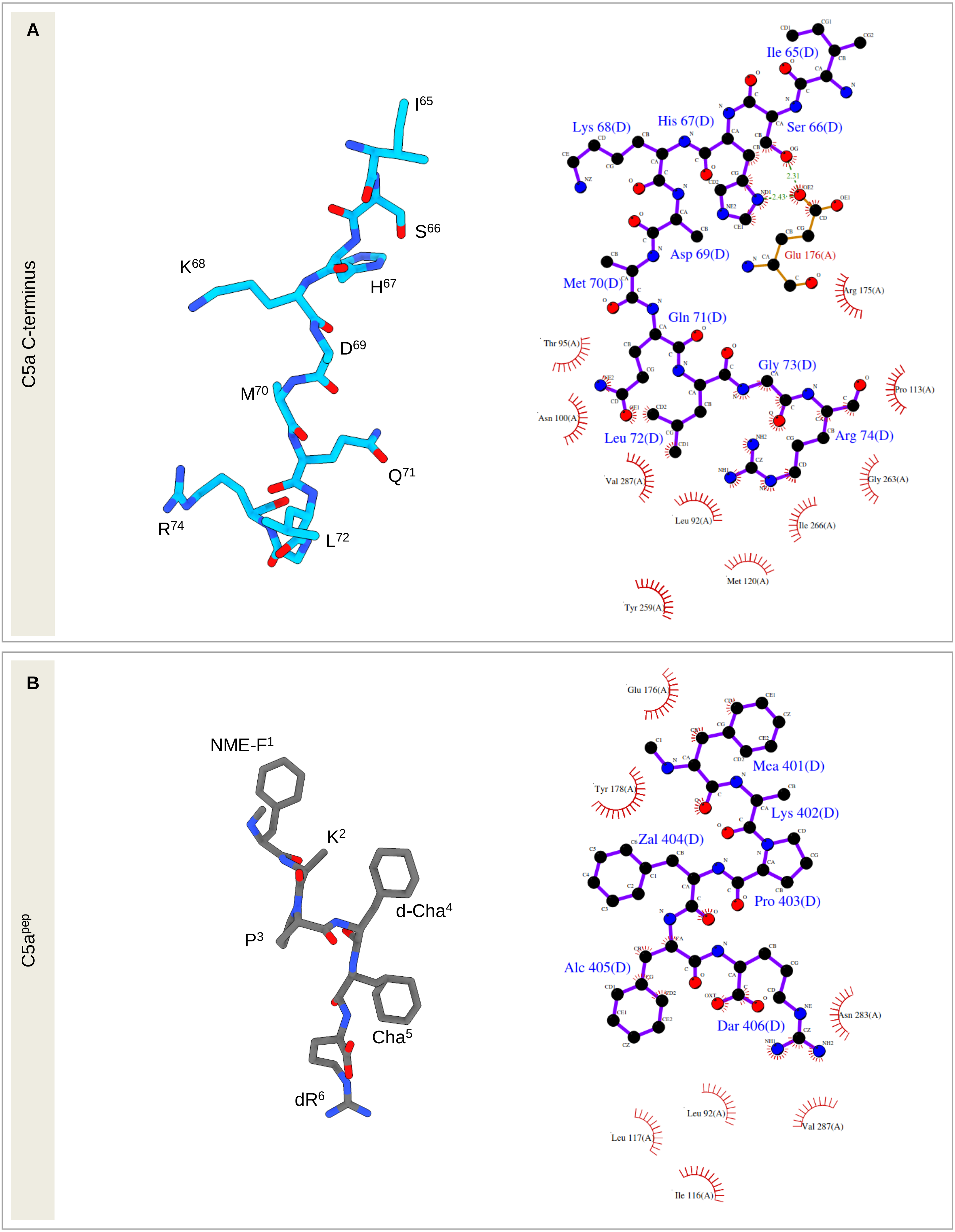

A
C5a C-terminus
I65
S66
K68
H67
D69
M70
Q71
R74
L72
Ile 65(D)
Lys 68(D)
His 67(D)
Ser 66(D)
2.31
2.43
Asp 69(D)
Glu 176(A)
Met 70(D)
Arg 175(A)
Gln 71(D)
Thr 95(A)
Gly 73(D)
Pro 113(A)
Asn 100(A)
Leu 72(D)
Arg 74(D)
Val 287(A)
Gly 263(A)
Leu 92(A)
Ile 266(A)
Met 120(A)
Tyr 259(A)
B
C5apep
NME-F1
K2
d-Cha4
P3
Cha5
dR6
Glu 176(A)
Mea 401(D)
Tyr 178(A)
Lys 402(D)
Zal 404(D)
Pro 403(D)
Alc 405(D)
Asn 283(A)
Dar 406(D)
Leu 92(A)
Val 287(A)
Leu 117(A)
Ile 116(A)

**Figure S12: Interaction interface of C5a C-terminus and C5a$^{pep}$ with C5aR1.**

**(A, B)** Overall atomic representation of C5a C-terminus and C5a$^{pep}$ (left) and residue contacts with C5aR1 (right). Interaction plots are generated using PDBSum (REF).

**A**

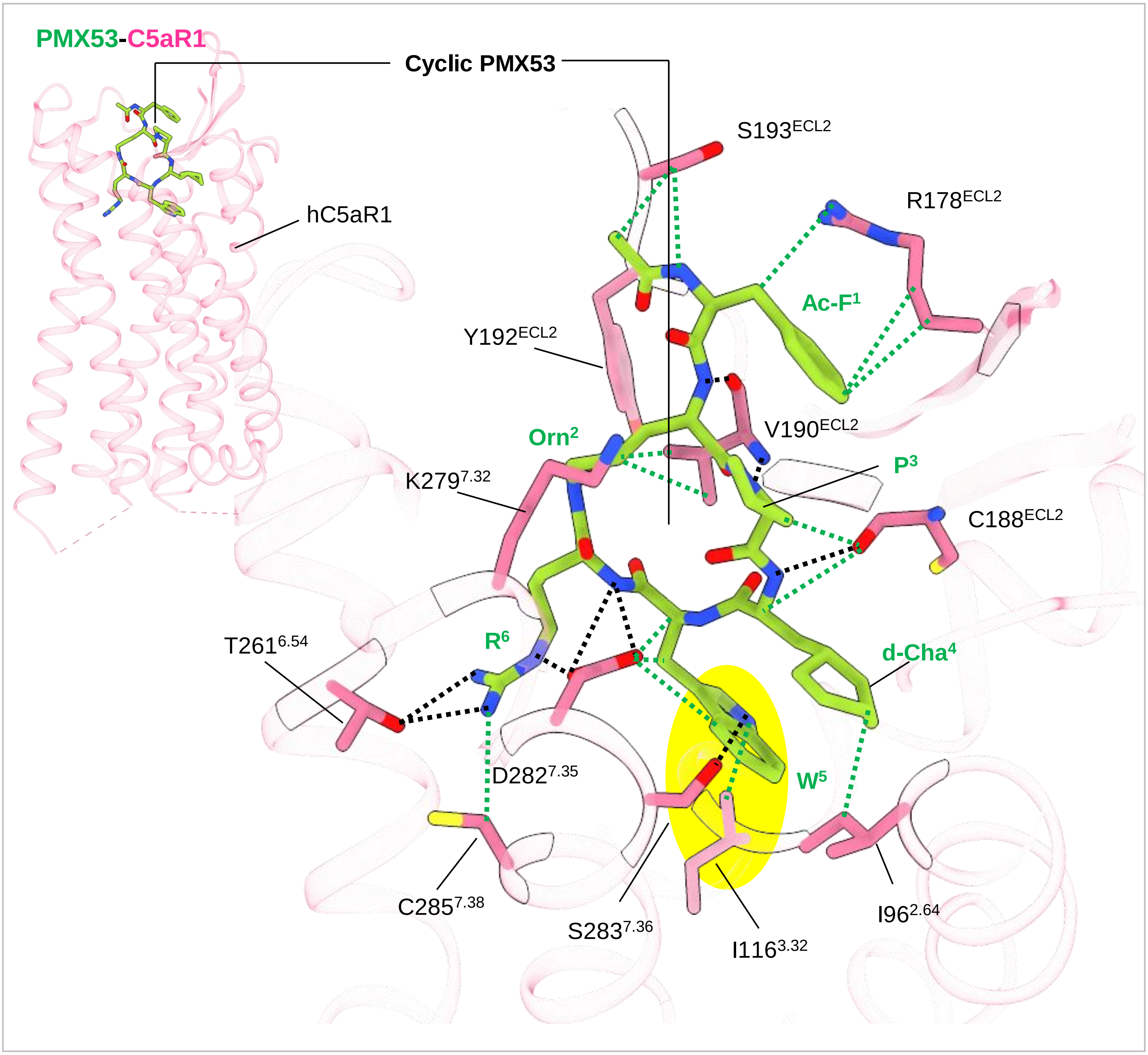
PMX53-C5aR1
Cyclic PMX53
hC5aR1
S193ECL2
R178ECL2
Ac-F1
Y192ECL2
Orn2
V190ECL2
P3
K2797.32
C188ECL2
T2616.54
R6
d-Cha4
D2827.35
W5
C2857.38
S2837.36
I1163.32
I962.64

**Figure S13: PMX53 engages with several extra residues of C5aR1 compared to C5a/C5a$^{pep}$.**

**(A)** Structure of PMX53 bound human C5aR1 (PDB: 6C1R). PMX53 makes crucial interactions with C5aR1 which are absent in the active state C5a/C5a$^{pep}$ bound C5aR1 structures. The bulky Trp$^{5}$ interacts with and blocks the movement of I116$^{3.32}$ reported to facilitate activation of C5aR1 (highlighted in yellow circle). (Polar interactions: Black dotted lines, Non-bonded contacts: Green dotted lines).

**A**

| C5aR1(C5a)-Go interface | | |
|---|---|---|
| **Chain A (C5aR1)** | **Distance (Å)** | **Chain B (Go)** |
| Asn71 (TM2) | Gly350 (3.79), Cys351 (3.84) | Gly350, Cys351 |
| Arg134 (TM3) | Cys351 (3.50), Leu353 (3.44) | Cys351, Leu353 |
| Leu137 (TM3) | Asn347 (3.08) | Asn347 |
| Pro141 (ICL2) | Ile343 (3.65), Ile344 (3.70) | Ile343, Ile344 |
| Ile142 (ICL2) | Leu195 (3.84) | Leu195 |
| Cys144 (ICL2) | Ile343 (3.78), Asn347 (3.77) | Ile343, Asn347 |
| Gln145 (ICL2) | Ile343 (3.56), Leu195 (3.68) | Ile343, Leu195 |
| Arg148 (ICL2) | Asn347 (3.31) | Asn347 |
| Arg233 (ICL3) | Asp341 (2.79) | Asp341 |
| Ala235 (ICL3) | Tyr320 (3.44), Asp341 (3.56) | Tyr320, Asp341 |
| Thr236 (ICL3) | Tyr354 (2.60), Ala345 (3.69) | Tyr354, Ala345 |
| Arg237 (ICL3) | Tyr354 (3.66) | Tyr354 |
| Ser238 (ICL3) | Tyr354 (3.48) | Tyr354 |
| Lys240 (TM6) | Leu353 (3.55), Tyr354 (3.23) | Leu353, Tyr354 |
| Thr241 (TM6) | Leu353 (3.14) | Leu353 |
| Val244 (TM6) | Leu353 (3.61) | Leu353 |
| Val245 (TM6) | Leu353(3.67) | Leu353 |
| Ala304 (TM7) | Tyr354 (2.85) | Tyr354 |

**B**

| C5aR1(C5a$^{pep}$)-Go interface | | |
|---|---|---|
| **Chain A (C5aR1)** | **Distance (Å)** | **Chain B (Go)** |
| Asn71 (TM2) | Gly350 (2.49) | Gly350 |
| Arg134 (TM2) | Cys351 (3.48) | Cys351 |
| Val138 (TM3) | Ile344 (3.76) | Ile344 |
| Pro141 (ICL2) | Ile344 (3.55) | Ile344 |
| Ile142 (ICL2) | Asn194 (3.43), Leu195 (3.78) | Asn194, Leu195 |
| Gln145 (ICL2) | Lys32 (3.80) | Lys32 |
| Lys146 (ICL2) | Lys32 (3.62) | Lys32 |
| Leu226 (TM5) | Leu353 (3.81) | Leu353 |
| Arg233 (ICL3) | Asp337 (3.87), Asp341 (2.58) | Asp337, Asp341 |
| Ala235 (ICL3) | Glu318 (3.22), Tyr320 (3.37) | Glu318, Tyr320 |
| Thr236 (ICL3) | Ala345 (3.58) | Ala345 |
| Ser238 (ICL3) | Tyr354 (2.77) | Tyr354 |
| Lys240 (TM6) | Tyr354 (3.11) | Tyr354 |
| Thr241 (TM6) | Leu353 (3.31) | Leu353 |
| Val244 (TM6) | Leu353 (3.48) | Leu353 |
| Val245 (TM6) | Leu353 (3.71) | Leu353 |
| Ala304 (TM6) | Gly352 (3.73) | Gly352 |

**Figure S14: (A, B)** List of C5a/C5a$^{pep}$-C5aR1 and Go interactions in the structures of C5a/C5a$^{pep}$-C5aR1-Go**.**
